# Structural basis of plp2-mediated cytoskeletal protein folding by TRiC/CCT

**DOI:** 10.1101/2022.07.25.501395

**Authors:** Wenyu Han, Mingliang Jin, Caixuan Liu, Qiaoyu Zhao, Shutian Wang, Yifan Wang, Yue Yin, Chao Peng, Yanxing Wang, Yao Cong

**Affiliations:** State Key Laboratory of Molecular Biology, National Center for Protein Science Shanghai, Shanghai Institute of Biochemistry and Cell Biology, Center for Excellence in Molecular Cell Science, Chinese Academy of Sciences, Shanghai 200031, China; University of Chinese Academy of Sciences, Beijing 100049, China; National Facility for Protein Science in Shanghai, Zhangjiang Lab, Shanghai Advanced Research Institute, CAS, Shanghai 201210, China

**Keywords:** Chaperonin TRiC/CCT, cochaperone plp2, tubulin/actin folding, ATPase cycle, cryo-EM

## Abstract

The eukaryotic chaperonin TRiC/CCT assists the folding of ~10% cytosolic proteins. The essential cytoskeletal proteins tubulin and actin are the obligate substrates of TRiC and their folding involves cochaperone and co-factors. Here, through cryo-EM analysis, we present a more complete picture of yeast TRiC-assisted tubulin and actin folding in the ATPase-cycle, under the coordination of cochaperone plp2. Our structures revealed that in the open C1 and C2 states, plp2 and substrates tubulin/actin engage with TRiC inside its chamber, one per ring. Noteworthy, we captured a ternary TRiC-plp2-tubulin complex in the closed C3 state, engaged with a full-length β-tubulin in the native folded state even loaded with a GTP, and with a plp2 occupying the opposite ring, not reported before. Another closed C4 state revealed an actin in the intermediate state of folding and a plp2 occupying the other ring. Intriguingly, along with TRiC ring closure, we captured a large translocation of plp2 within TRiC chamber coordinating substrate translocation on the CCT6 hemisphere, potentially facilitating substrate stabilization and folding. Our findings provide structural insights into the folding mechanism of the major cytoskeletal proteins tubulin/actin under the coordination of the complex biogenesis machinery TRiC and plp2, and could extend our understanding on the links between cytoskeletal proteostasis and related human diseases.

## Introduction

The eukaryotic group II chaperonin TRiC/CCT assists the folding of about 10% cytosolic proteins, including many key structural and regulatory proteins. TRiC plays an essential role in maintaining cellular protein homeostasis, and dysfunction of TRiC is closely related to cancer and neurodegenerative diseases (*1–3*). The key cytoskeletal proteins actin and tubulin, the most abundant and highly conserved proteins in eukaryotic cells, are obligate substrates of TRiC (*4–7*). This chaperonin is also essential for the folding of cell cycle regulator CDC20 (*8*), and many proteins involved in oncogenesis, such as p53, VHL tumor suppressor, and STAT3 (*9–11*). TRiC undergoes large conformational changes to encapsulate and release bound substrate proteins during its ATP-dependent folding cycle (*12–14*).

TRiC is the most complex chaperonin system, consisting of two octameric rings that stacked back-to-back. Each ring of TRiC encompasses eight paralogous subunits, CCT1-CCT8 (sharing ~23%-35% sequence identity for yeast TRiC), arranged in a specific order (*15–20*). Corroborate to this, TRiC displays subunit specificity in ATP consumption, ring closure, and substrate recognition and folding (*15, 18, 21*). Each TRiC subunit consists of an equatorial domain (E domain) that contains the ATP-binding site and forms the intra- and inter-ring contacts, an apical domain (A domain) that comprises the interaction sites with target proteins, and an intermediate domain (I domain) that connects the other two domains (*22–24*).

The filament-forming actin and tubulin are abundant cytoskeletal proteins that support diverse cellular processes. Owing to the unique properties of these proteins, a remarkably complex cellular machinery consisting minimally of the chaperonin TRiC, prefoldin (PFD), phosducin-like proteins, and tubulin cofactors has evolved to facilitate their biogenesis (*25*). TRiC cooperates with co-chaperone PFD in the folding of actins and tubulins (*26, 27*), in which process PFD stabilizes substrate proteins and delivers them to the open TRiC cavity (*25, 26, 28–31*). Phosducin-like protein, including three homologue families (PhLP1, PhLP2, and PhLP3), shares an N-terminal helical domain, a central thioredoxin fold domain (TRDX domain), and a C-terminal extension (*32, 33*). It has been shown that PhLP2 is essential for ciliogenesis and microtubule assembly (*34, 35*), and temperature sensitive alles of plp2, the homologue of PhLP2 in *S. cerevisiae,* caused defective in TRiC-mediated actin and tubullin function (*35*). Mammalian PhLP2A modulates the TRiC chaperonin activity during folding of cytoskeletal proteins, and yeast plp2 considerably accelerates actin folding rate than that assisted by TRiC alone (*36–38*). It has been indicated that the ciliary precursor tubulin needs to be folded by TRiC with assistance of PhLP2 (*34*). On the basis of previous structural studies on TRiC-actin and TRiC-tubulin complexes at intermediate or relatively low resolutions (*6, 7, 39–42*), a recent study reported cryo-EM structures of human TRiC-PhLP2A-actin and TRiC-tubulin both in the closed state at near atomic resolution, and an open TRiC structure with potential substrate density located in the inter-ring septum (*43*). Still, how PhLP2 coordinates with the TRiC-assisted tubulin/actin folding, especially the role of PhLP2 in TRiC-assisted tubulin folding, in the undergoing ATP-driven TRiC conformational cycle remains to be further explored.

In the present study, we present an ensemble of cryo-EM structures of *S. cerevisiae* TRiC along its ATPase cycle, with simultaneously engaged plp2 and substrate actin or tubulin inside its chamber, one per ring, in the resolution range of 3.05 Å to 4.55 Å. In the open C1 and C2 states, we observed plp2, with detectable TRDX domain, contacting the CCT3/1/4 subunits of TRiC and the substrate density in opposite ring mostly located in the CCT6 hemisphere. Our structural analyses further revealed the upward translocation of both plp2 and substrates alone with TRiC ring closure, and the molecular details of the unique TRiC-plp2 interaction mode. Strikingly, we captured the TRiC-plp2-tubulin tertiary complex, with tubulin reached the fully folded state and loaded with a GTP since its “born”, not observed before. Collectively, our study suggests that plp2 serves a cochaperone of TRiC-assisted substrate tubulin/actin folding, and reveals stepwise substrate folding process within TRiC chamber accompanying its ATP-driven conformational cycle, providing mechanistic insights on how plp2 participates in the substrate tubulin/actin folding process assisted by TRiC.

## Results

### Simultaneous binding of plp2 and substrates inside open-state TRiC chamber

To capture a complete picture of TRiC-assisted substrate folding and to reveal the role of plp2 in this process, we purified *S. cerevisiae* TRiC, plp2, and actin, respectively (Fig. S1A). We then incubated them together without adding nucleotide, which were confirmed to form a complex by native gel analysis (Fig. S1B). Our mass spectrometry result further suggested that TRiC co-exists with actin and plp2, in addition to endogenous yeast β-tubulin, which could be co-purified with TRiC from yeast in a lesser amount (Table S2). Here we followed a previous report to pretreat actin and plp2 with EDTA that could unfold actin (*37*). Under this condition, actin was indeed unfolded while plp2 not, suggested by the thermo-stability assay (Fig. S1C-F).

Our cryo-EM study on such system revealed an open-state TRiC map loaded with two extra densities inside its chamber, one in each ring, at the resolution of 4.55Å (termed “C1” state, Fig. 1A-B, S2A-C, Table S1). The two densities appeared distinct (Fig. 1B). Overall, the TRiC conformation is comparable to our previous yeast TRiC structure in the nucleotide partially preloaded (NPP) state (Fig. S2D) (*18*), both displaying the following characteristic features: (1) CCT1 being the most outward tilted subunit (Fig. 1A), a feature common in yeast, bovine, and human open-state TRiC (*12, 18, 39, 44*); (2) the A and I domains of CCT2 as a whole displaying a bent feature (Fig. 1A), unique for yeast TRiC-NPP state (*18*); (3) CCT7 also being slightly outward tilted (Fig. S2D). These features allowed us to assign the TRiC subunits and build an atomic model for the C1 map (Fig. S2E). In this map, CCT3/6/8 subunits also bear pre-loaded nucleotides as in the free TRiC-NPP map (Fig. S2F) (*18*). Collectively, these data indicate that the C1 TRiC is in the NPP state.

Further inspection of the C1 map revealed that in the cis-ring chamber, the extra density attaches to the consecutive CCT3/1/4 subunits, including the A domain of CCT3/4 and the E domain of CCT3/1 (Fig. 1C). The lower portion of the extra density matches well with the TRDX domain of yeast plp2 (Fig. 1E), indicating the cis-ring extra density most likely belongs to plp2. Corroborate to this, our chemical crosslinking and mass spectrometry (XL-MS) analysis also detected XLs between plp2 and CCT3/1/4 (Fig. 1G, C, Table S3). Regarding the extra density in the trans-ring chamber, it attaches to the CCT8/6/3 subunits, involving the A domain of CCT6/3 and the E domain of all the three subunits (Fig. 1D, F). Still, this extra density is less well resolved, and hardly to judge it belongs to actin or tubulin. Moreover, our XL-MS analysis revealed a XL between actin and CCT6 (Fig. 1G, D, Table S3). Combined with our mass spectrometry result, indicating the co-exitance of endogenous tubulin with TRiC although in a lesser amount (Table S2), we then postulate that the trans-ring extra density may correlate to the bound substrates actin and tubulin, which are in the initial stage of folding and remain very dynamic. Therefore, our C1 map suggests that in the open NPP state, yeast TRiC can simultaneously engage with plp2 in one ring and the essential cytoskeletal proteins actin or tubulin in the opposite ring. This is distinct from the recent human open state TRiC structure, with potential substrate (tubulin/actin) density in the inter-ring septum (*43*).

### Plp2 remains in the same location after ATP binding to TRiC

To capture a thorough picture of plp2-mediated tubulin/actin folding assisted by TRiC conjugated with its ATPase-cycle, we incubated the above-mentioned system with ATP-AlFx, a nucleotide analog has been used to mimic ATP-hydrolysis-transition state in previous structural studies on TRiC and TRiC-substrate complexes (*12, 19, 28, 39, 43, 45, 46*). Our cryo-EM study on this system revealed two conformational states of TRiC, one in the open state (Fig. S3A), and the other in the closed state (discussed later, Fig. S3A). The open and closed state of TRiC displayed a population distribution of 27% and 73%, respectively (Fig. S3A-B). This population distribution is very distinct from that of our previous substrate-free yeast TRiC in the same chemical condition, in which almost all TRiC complexes are in the closed conformation (*19*). Moreover, our NADH-coupled assay suggested that the ATPase activity of TRiC remains the same regardless of the presence or absence of plp2 and actin (Fig. S5). Taken together, the presence of plp2 and actin does not alter the TRiC ATPase activity, instead, it may tune the allosteric network of TRiC, with 27% of TRiC particles delayed transferring to the closed state (more in discussion).

Further focused classification of the TRiC-open dataset led to an open state map, with TRiC engaged with two extra densities, one inside each chamber (termed “C2” state, Fig. 2A-B, S3A, C-D, S4A-B). In the C2 map, all the TRiC subunits load with nucleotide densities that match ATP reasonably well (Fig. S4D). Also, the overall open TRiC conformation resembles that of TRiC-AMP-PNP (Fig. S4C) (*18*), with the A-I domains of CCT2 unbent and CCT7 tilt inwards, making the complex less asymmetric and stabilized slightly. Altogether, these data suggest that the C2 map represents TRiC in the ATP-bound state. Further inspection of this map showed that the cis-ring extra density attaches to the A domain of CCT3/4 and the E domain of CCT3/1, and this density matches the TRDX domain of plp2 well (Fig. 2C-D), overall similar to the plp2 observed in the C1 map (Fig. 1C, E). The trans-ring extra density appears distinct in size and shape to that in the cis-ring, and most likely represents substrate tubulin/actin (Fig. 2E-F). It forms contacts with CCT3/1/4 subunits, implying a potential substrate translocation upon ATP binding to TRiC. Taken together, our C1 and C2 maps both suggest that, in either the NPP or the ATP-bound open-state yeast TRiC, plp2 and substrate actin or tubulin can simultaneously occupy TRiC chamber, one per ring, not seen in the open state human TRiC structure (*43*).

### TRiC cooperates with plp2 in the folding of tubulin/actin accompanying TRiC ring closure

For the closed-state TRiC dataset of the ATP-AlFx incubated sample, we performed further focused 3D classifications on the inner chamber of each of the two TRiC rings, and obtained three cryo-EM maps, termed C3, C4, and C5 state at the resolution of 3.39, 3.20, and 3.05 Å, respectively (Fig. S3A, C, E). Specifically, for the C3 and C4 maps, the two extra densities, one inside each chamber, appear in distinct shape (Fig. 3A-B, 4A-B); while for the C5 map, it displays two almost identical extra densities, one in each ring (Fig. 5A-B). For the closed TRiC maps, a characteristic kink feature in the apical domain α-helical protrusion of one subunit (Fig. S4E), corresponding to the unique feature of CCT6 (*15*), and a protruding density corresponding to the inserted CBP-tag on the A domain of CCT3 could be directly visualized (Fig. S4F). These characteristics allowed us to identify the subunits and build a model for each of the C3, C4, and C5 maps (Fig. S4G-L).

**Fig. 1.**
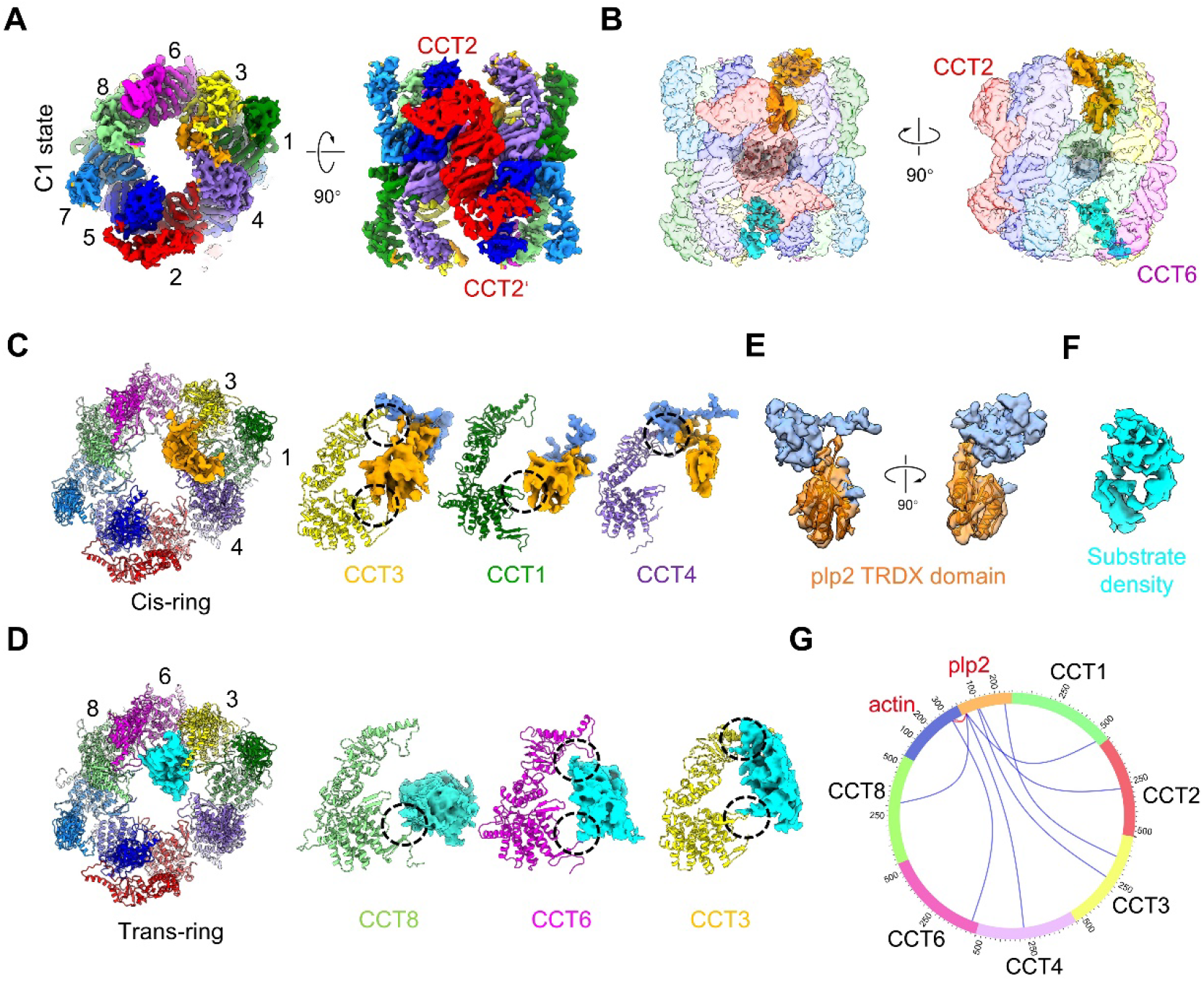
Cryo-EM structure of TRiC-plp2-substrate showed plp2 and substrate existed within distinct TRiC chamber at TRiC-NPP state. (A) Top and side view of TRiC-plp2-substrate map at C1 state, with CCT1-CCT8 showing in green, red, yellow, medium purple, blue, magenta, dodger blue and light green, respectively, which color scheme of TRiC is followed throughout. (B) Side view of C1 state with unsharpened transparent map to show the location of extra density (cis-ring in orange and trans-ring in cyan) and unstructured tail mass (grey) relative to TRiC chamber. (C) Top view of C1 model showed the extra density map in cis-ring interacted with CCT3/1/4, which were indicated by black circle. (D) Bottom view of C1 model showed that extra density in trans-ring interacted with CCT8/6/3, which were indicated by black circle. (E) The lower portion of extra density in cis-ring fitted well with the TRDX domain (orange) of plp2, with the relative dynamic N domain (cornflower blue) floating. (F) The extra density in trans-ring appeared poor structural features and connectivity. (G) The circular plot of XL-MS results of TRiC-plp2-actin, which showed there were 6 unique XLs between CCT1/2/3/4/8 and plp2, while there was only one unique XL between CCT6 and actin. There were also 2 XLs between actin and plp2 (red lines). However, there are no XLs about tubulin.

In the C3 map, the cis-ring extra density matches the plp2 model well, with the TRDX domain and about half of the N-terminal helical domain and C-terminal extension being resolved (Fig. 3C-D). It appeared that the plp2 lies underneath the dome, interacting with the CCT6/3/1/7/5/2 subunits of TRiC (Fig. 3B-C, detailed TRiC-plp2 interactions will be presented later). Collectively, our structures suggested that accompanying TRiC ring closure, the plp2 TRDX domain displays obvious translocation from the CCT6 hemisphere (in the open C1 and C2 states) to the CCT2 hemisphere (in the closed C3 state, Fig. 1C, 2C and 3C). In addition, the trans-ring extra density matches the structural features of full-length folded tubulin very well (Fig. 3C, E-G). Noteworthy, this TRiC-plp2-tubulin ternary structure has not been captured before, although being detected biochemically (*35*). Conjugating with TRiC ring closure and plp2 translocation, tubulin also upward shifted and appeared stabilized, although remained on the CCT6 hemisphere as in the open C1/C2 states. Also, the folded tubulin density exhibits structural features of the S355-A364 loop, usually used to distinguish β-tubulin from α-tubulin (*47, 48*), confirmed the resolved tubulin is indeed β-tubulin (Fig. 3F, H), in line with our mass spectrometry result (Table S2). Surprisingly, this folded β-tubulin also revealed a bound nucleotide density matched GTP very well (Fig. 3H), indicating that the newly made β-tubulin already loaded with a GTP before released from TRiC chamber. Inspection of the interaction between TRiC and tubulin revealed that the N- and C-domain of tubulin attached to the A- and Edomain of CCT8/6/3/1 subunits, mainly with the Helix11 of CCT8/6/3, loop^H11^ of CCT8/6/3/1, loop^H10^ of CCT6, the protrusion loop of CCT8/6/1, and the E-domain stem loop of CCT6/3/1 subunits (Fig. 3I, S4N, Table S4). Further inspection of the surface property showed that the abovementioned interaction footprint on TRiC is more positively charged, compensate the overall negatively charged tubulin in the interaction interface (Fig. 3J), and tubulin also interacted with TRiC through hydrophilic interaction (Fig. 3K). Collectively, our yeast TRiC-plp2-tubulin structure enabled us to capture unexpected features compared with previous TRiC-tubulin structures (Kelly et al., 2022; Munoz et al., 2011), including a plp2 molecule co-existed with substrate tubulin in TRiC chamber (one per ring), a bound GTP in the newly made β-tubulin, and the fully folded tubulin to its native state, whereas the tubulin I domain remains disordered in the human TRiC-tubulin structure in the absence of plp2 (*43*). Therefore, the presence of plp2 could play a role in stabilizing and folding the substrate tubulin.

**Fig. 2.**
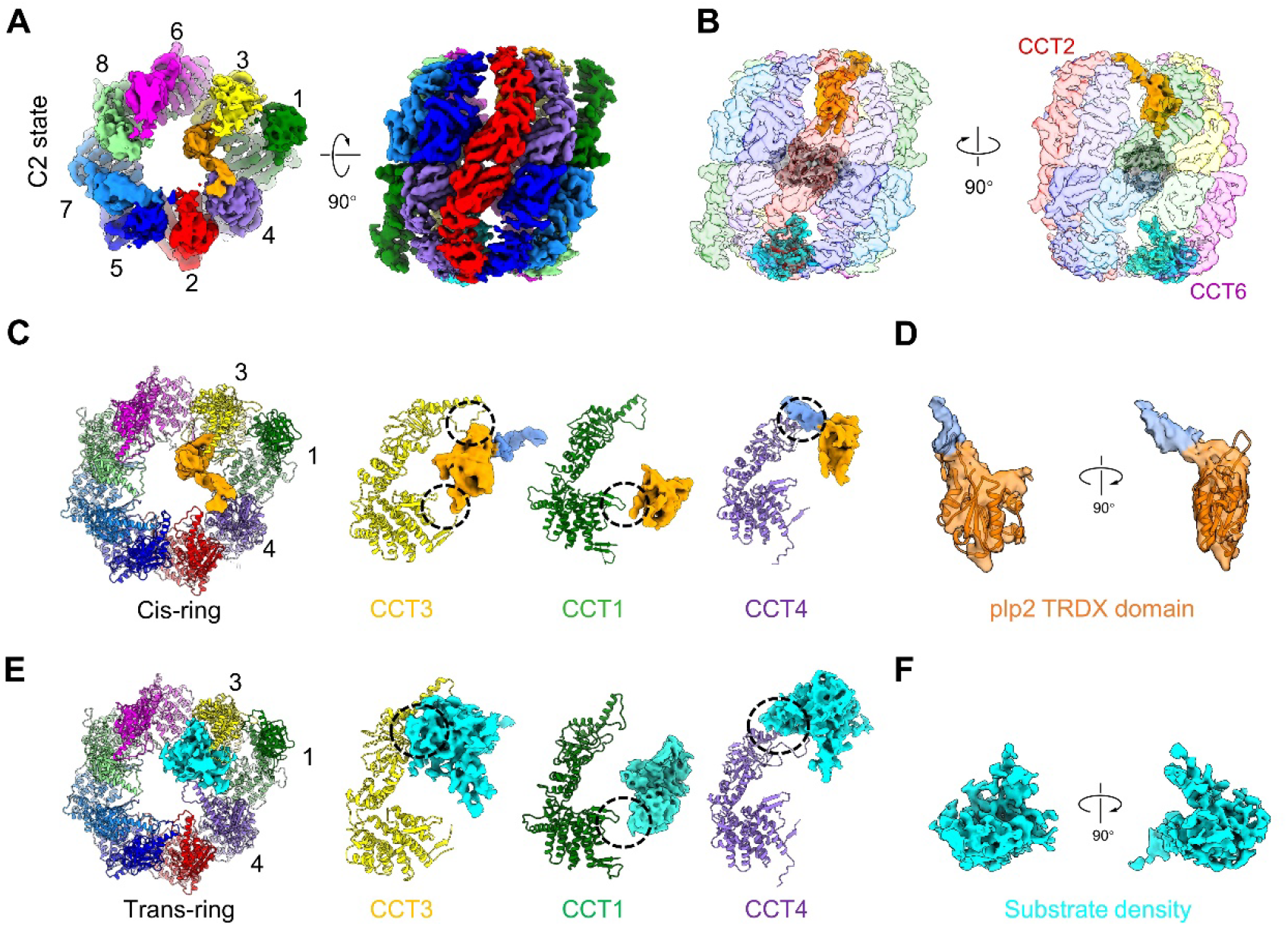
Cryo-EM structure of TRiC-plp2-substrate with ATP-AlFx at open state. (A) Top and side view of TRiC-plp2-substrate with ATP-AlFx at open C2 state. (B) Side view of C2 with unsharpened transparent map to show the location of extra density (cis-ring in orange and trans-ring in cyan) and unstructured tail mass (grey) relative to TRiC chamber. (C) Top view of C2 model showed the extra density map in cis-ring interacted with CCT3/1/4, which was indicated by black cycle and consistent with C1. (D) The lower portion of extra density in cis-ring of C2 also fitted well with the TRDX domain (orange) of plp2, with the relative dynamic N domain (cornflower blue) floating. (E) Bottom view of C2 model showed the extra density map in trans-ring also interacted with CCT3/1/4, which were indicated by black circle. (F) The extra density in transring of C2 still exhibited poor structural features and connectivity as C1 map.

**Fig. 3.**
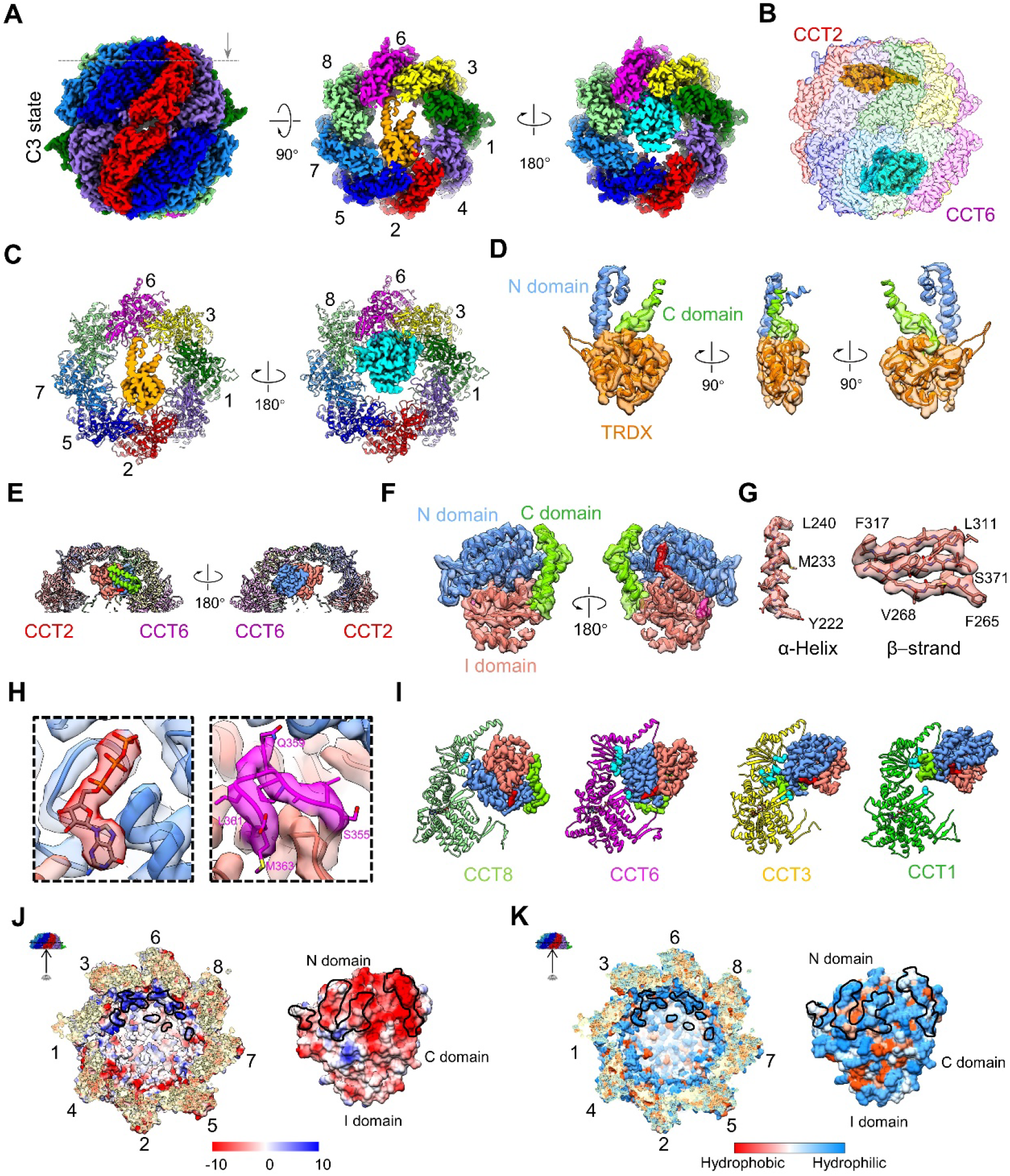
Both plp2 and β-tubulin existed within distinct closed TRiC chamber with GTP bound tubulin since its “born”. (A) Side view (left), top view (middle) and bottom view (right) of closed C3 state. The grey dashed line was the clipped orientation for top and bottom view to better show the extra density (cis-ring in orange and trans-ring in cyan) were located underneath the dome of TRiC. (B) Side view of C3 showed the location of extra densities relative to TRiC chamber. (C) Top and bottom view showed that the extra density in cis-ring interacted with CCT6/3/1/2/5/7 subunits and the extra density in trans-ring interacted with CCT8/6/3/1 subunits. (D) The extra density in cis-ring of C3 fitted well with the plp2 model, with N domain showing in cornflower blue, TRDX domain in orange and C domain in chartreuse, which color scheme of plp2 is followed throughout. (E) Different side view showed substrate density in trans-ring mainly occupied CCT6 hemisphere. (F) Extra density in trans-ring of C3 fitted well with β-tubulin model (PDB:5W3F), N-, I- and C-domain of tubulin are colored in deep sky blue, salmon and green, respectively, which the color scheme of tubulin is followed throughout. (G) Resolved high-resolution structural features of tubulin I domain, the relative dynamic domain which had poor connectivity in previous TRiC-tubulin structure. (H) Magnified views to show the GTP model fitted well with our density and the short S355-A364 loop to distinguish α- or β-tubulin. (I) Interaction interface analysis by PISA showed that tubulin attached with the A- or E-domain of CCT8/6/3/1. The residue of TRiC subunits in proximity to tubulin (less than 4 Å) are shown as cyan balls (Table S4). (J-K) Interaction interface analysis of surface properties between TRiC and β-tubulin, showing β-tubulin interacting with TRiC through electrostatic (J) and hydrophilic (K) interactions.

In the C4 map, the cis-ring extra density was also identified as plp2, displaying the same conformation and binding location as that in C3 (Fig. 4C-D, S4M). For the trans-ring extra density, it mainly occupies CCT6 hemisphere, as that in the open C1 and C2 states. This density overall matches the actin model although being less well resolved, indicating a relative dynamic status of actin potentially in the intermediate state of folding (Fig. 4E-F). It appeared that actin mainly attaches to CCT8/6/3/1, with the unstructured portion stretching into the chamber (Fig. 4E). Our C4 map is overall similar to the recent structure of human TRiC-actin-PhLP2A also in the closed state (*43*), whilst here actin is more in the intermediate status of folding (more in discussion). **Interactions between plp2 and closed state TRiC**

**Fig. 4.**
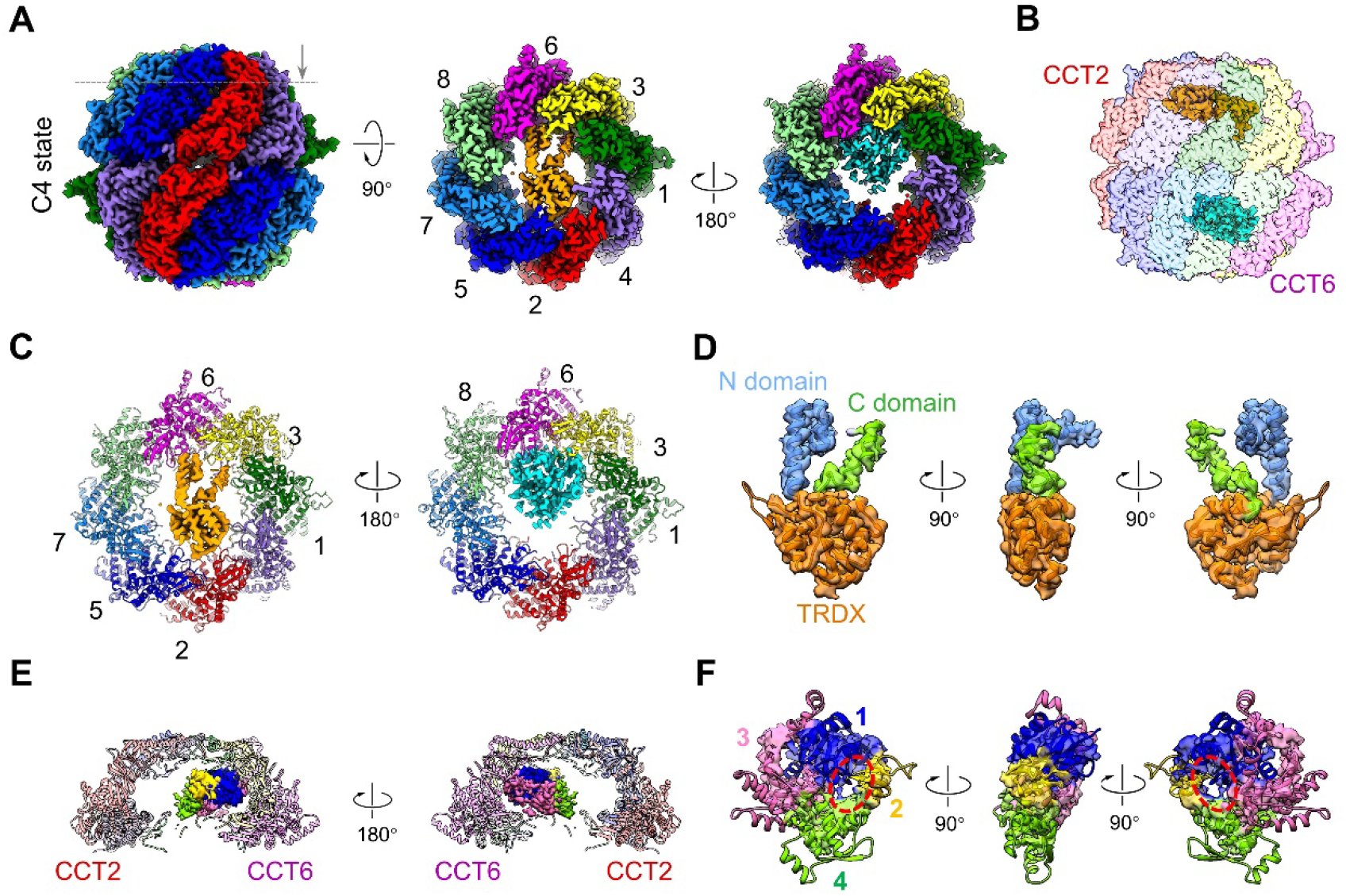
Both plp2 and actin existed within distinct closed TRiC chamber. (A) Side view (left), top view (middle) and bottom view (right) of closed C4 state. The grey dashed line was the clipped orientation for top and bottom view to better show the extra density (cis-ring in orange and trans-ring in cyan) were lied across the chamber. (B) Side view of C4 showed the location of extra densities relative to TRiC chamber. (C) Top and bottom view showed that the extra density in cis-ring interacted with CCT6/3/1/2/5/7 subunits and the extra density in trans-ring interacted with CCT8/6/3/1 subunits. (D) The extra density in cis-ring of C4 fitted well with the plp2 model. (E) Different side view showed substrate density in trans-ring mainly occupied CCT6 hemisphere with unstably portion tilting toward CCT2 hemisphere. (F) Rigid body fitting of extra density in trans-ring of C4 map with actin model (PDB:1YAG), subdomain 1-4 is colored in blue, gold, hot pink and green, respectively, which the color scheme of actin is followed throughout. Red cycle indicated the cleft between subdomain 2 and subdomain 4 in native actin.

Regarding the C5 map resolved at the resolution of 3.05 Å, the plp2 model fits in the extra densities in both rings very well (Fig. 5C-E, S3E), indicating plp2 could occupy both chambers of TRiC, potentially depending on the concentration of plp2. Moreover, in the cis-ring of C3/C4 and both rings of C5 map, the engaged plp2 exhibits similar architecture and orientation within closed TRiC chamber (Fig. S4M), thus we used the better resolved cis-ring plp2 in C5 for subsequent structural analysis. Here, other than the TRDX domain, several α-helixes of the N-terminal helical domain and part of its flexible C-terminal extension were well resolved (Fig. 5D-E). Specifically, the N-terminal helical domain and the C-terminal extension of plp2 attaches to CCT6/3/1 subunits from the CCT6 hemisphere, and the TRDX domain of plp2, originally engaged with CCT3/1/4 subunits in the open C1/C2 states, now translocated and engaged with CCT2/5/7 from the CCT2 hemisphere across the inner chamber (Fig. 5C, F). Collectively, our data suggest that both plp2 and cytoskeletal substrates tubulin/actin translocate within TRiC chamber accompanying TRiC-ring closure, with plp2 stretching across the inner chamber of TRiC, while tubulin and actin interacting mainly with the CCT6 hemisphere subunits.

**Fig. 5.**
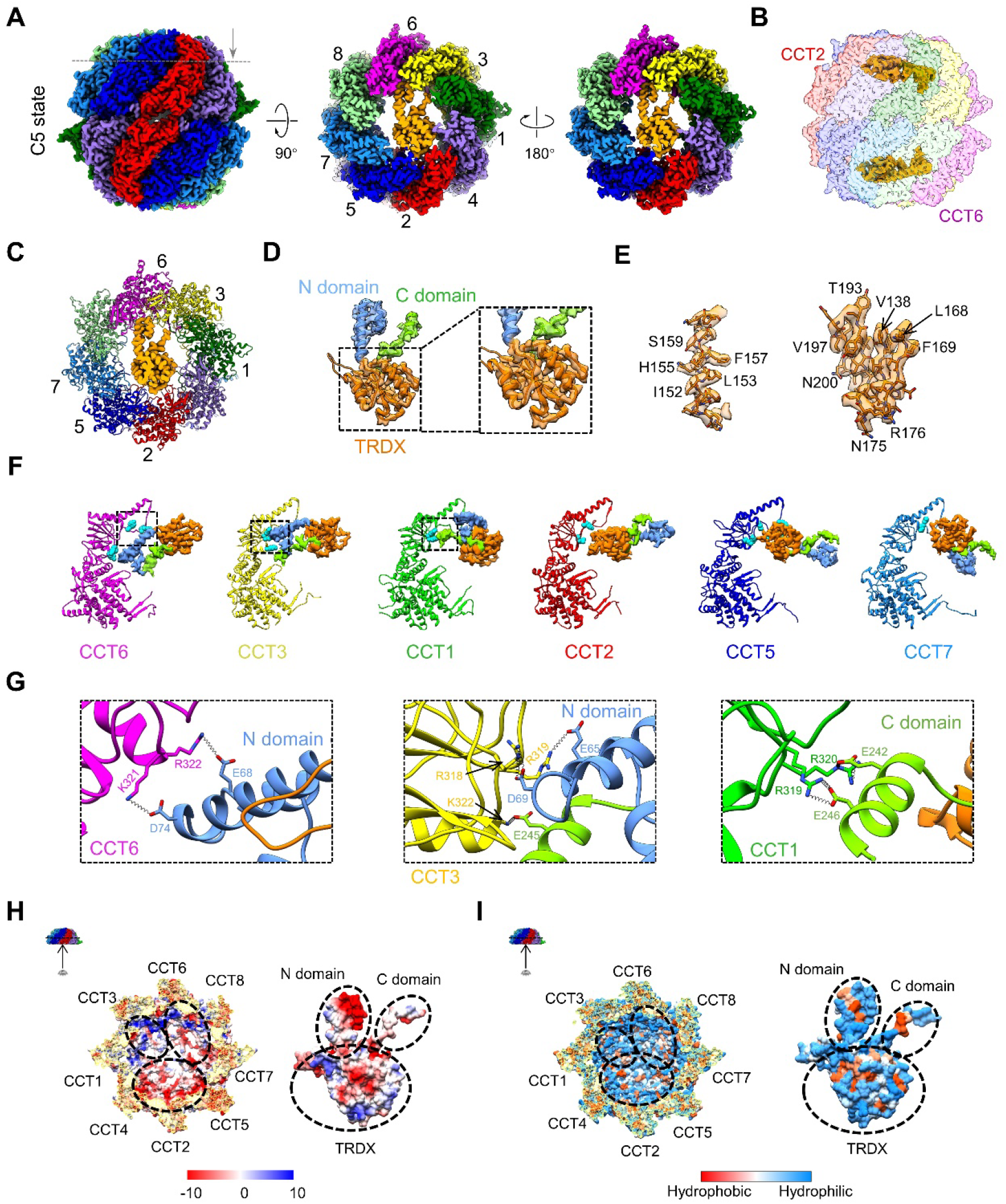
The binding motif between TRiC and plp2 in closed TRiC. (A) Side view (left), top view (middle) and bottom view (right) of closed C5 state. The grey dashed line was the clipped orientation for top and bottom view to better show the extra density were located underneath the dome of TRiC. (B) Side view of C5 showed the location of extra densities relative to TRiC chamber. (C) Top view showed that plp2 density interacted with CCT6/3/1/2/5/7 subunits. (D) Extra densities in cis- and trans-ring of C5 fitted well with plp2 model. Here, other than the TRDX domain of plp2, several α-helixes of the N-terminal helix domain, and part of its flexible C-terminal extension were well resolved. (E) Resolved high-resolution structural features of plp2 in C5. (F) Interaction interface analysis by PISA showed that plp2 attached with the A domain of CCT6/3/1/2/5/7. The residue of TRiC subunits in proximity to plp2 (less than 4 Å) are shown as cyan balls (Table S5). (G) Magnified views of the region in black box of (F), indicated with black dotted spring to show the salt bridges formed between plp2 and CCT6/3/1 subunits. (H-I) Surface analysis of coulombic and hydrophobic interaction were shown respectively. Plp2 mainly interacted with TRiC subunits through electrostatic interaction at closed state, which also had polar interface as a supplementary.

Our further structural analysis revealed salt bridges and hydrogen bonds (H-bound) formed mainly between the A-domain of CCT6/3/1/2/5/7 and the corresponding plp2 structural elements (Fig. 5F-G, Table. S5). Moreover, inspection of the electrostatic interaction between plp2 and TRiC revealed that the more positively charged CCT6 hemisphere is complementary with the plp2 N-terminal helical domain, and the more negatively charged CCT2 hemisphere complementary with the lower edge of the plp2 TRDX domain (Fig. 5H). In addition, plp2 and TRiC both display hydrophilic properties in their interaction interface (Fig. 5I). Taken together, TRiC and plp2 interact through a combination of electrostatic and hydrophilic interactions.

Moreover, as a control experiment, we performed further cryo-EM study on yeast TRiC-actin in the presence of ATP-AlFx but in the absence of plp2. We found that there is no detectable extra density inside closed TRiC chamber (Fig. S6), indicating no engaged substrate in TRiC in the absence of plp2. This is in line with a previous study on bovine TRiC-actin, which suggested that actin may be very dynamic and cannot be resolved even within the closed TRiC chamber (*39*). Taken together, plp2 may serve as a cochaperone facilitating substrate stabilization and folding within TRiC chamber.

## Discussion

Chaperonin TRiC and the phosducin-like proteins play critical roles in mediating the biogenesis of major cytoskeletal proteins actin and tubulin, thus with their activity being linked to any cellular process that depends on the integrity of the microfilament and microtubule-based cytoskeletal systems (*25, 49*). Still, how PhLP2 coordinates with TRiC in the folding of cytoskeletal proteins, especially tubulin, accompanying ATP-driven TRiC ring closure remains to be further explored. Here, we present ternary yeast TRiC-plp2-tubulin and TRiC-plp2-actin structures in the closed state (Fig. 3, 4), with the first structure not reported before; and also TRiC-plp2-substrate structures in the open NPP (C1) and ATP-bound (C2) states, with plp2 and substrate occupying different chambers (Fig. 1, 2), distinct from the recent report (*43*). Strikingly, our TRiC-plp2-tubulin (C3) structure suggested that with the assistant of plp2, yeast β-tubulin reaches the fully folded state with an ordered I domain and even loaded with a GTP since its “born” before released from TRiC chamber (Fig. 3). Our structural analyses combined with biochemical and mass spectrometry data depict a more complete picture of the plp2-mediated folding of major cytoskeletal proteins tubulin/actin by TRiC alone with its conformational cycle, reveal the role of plp2 potentially as a cochaperone in stabilization and folding of substrates tubulin/actin within TRiC chamber. Our study sheds lights on the structural basis of biogenesis of tubulin and actin mediated by TRiC cooperated with plp2.

### Mechanism of plp2 and TRiC coordinated tubulin/actin folding accompanying TRiC ring closure

Based on our results, we put forward a mechanism of plp2-mediated tubulin/actin folding by TRiC accompanying its ATPase cycle (Fig. 6). First, free NPP state TRiC can spontaneously engage with plp2 and substrates tubulin or actin in their initial folding state, one per ring, forming the C1 state (Step 1). In this stage, the plp2 TRDX domain engages with CCT3/1/4 subunits, and substrate mainly interacts with CCT8/6/3 on the CCT6 hemisphere. Subsequently, ATP binding makes TRiC more symmetrical and stabilized, and plp2 remains in the original location, forming the C2 state (Step 2). Afterwards, accompanying TRiC ring closure driven by ATP-hydrolysis, both plp2 and substrates tubulin/actin translocate upwards to lie underneath the dome, with plp2 stretching across the TRiC chamber, forming C3/C4 states (Step 3). With TRiC ring closure, the complex becomes more symmetrical and the generated mechanical force could assist substrate to overcome the energy barrier towards correct folding. Indeed, in this state, the substrate is better structured especially for tubulin with the assistance of plp2. Eventually, after TRiC releasing γ-phosphate and reopening its chamber, we postulate substrate and plp2 could be released from the chamber (Step 4). After this, plp2 could be reused by TRiC, and for the released not fully folded substrate (such as actin), they may need multiple rounds of binding and folding within TRiC to reach it native folded state (*50*).

**Fig. 6.**
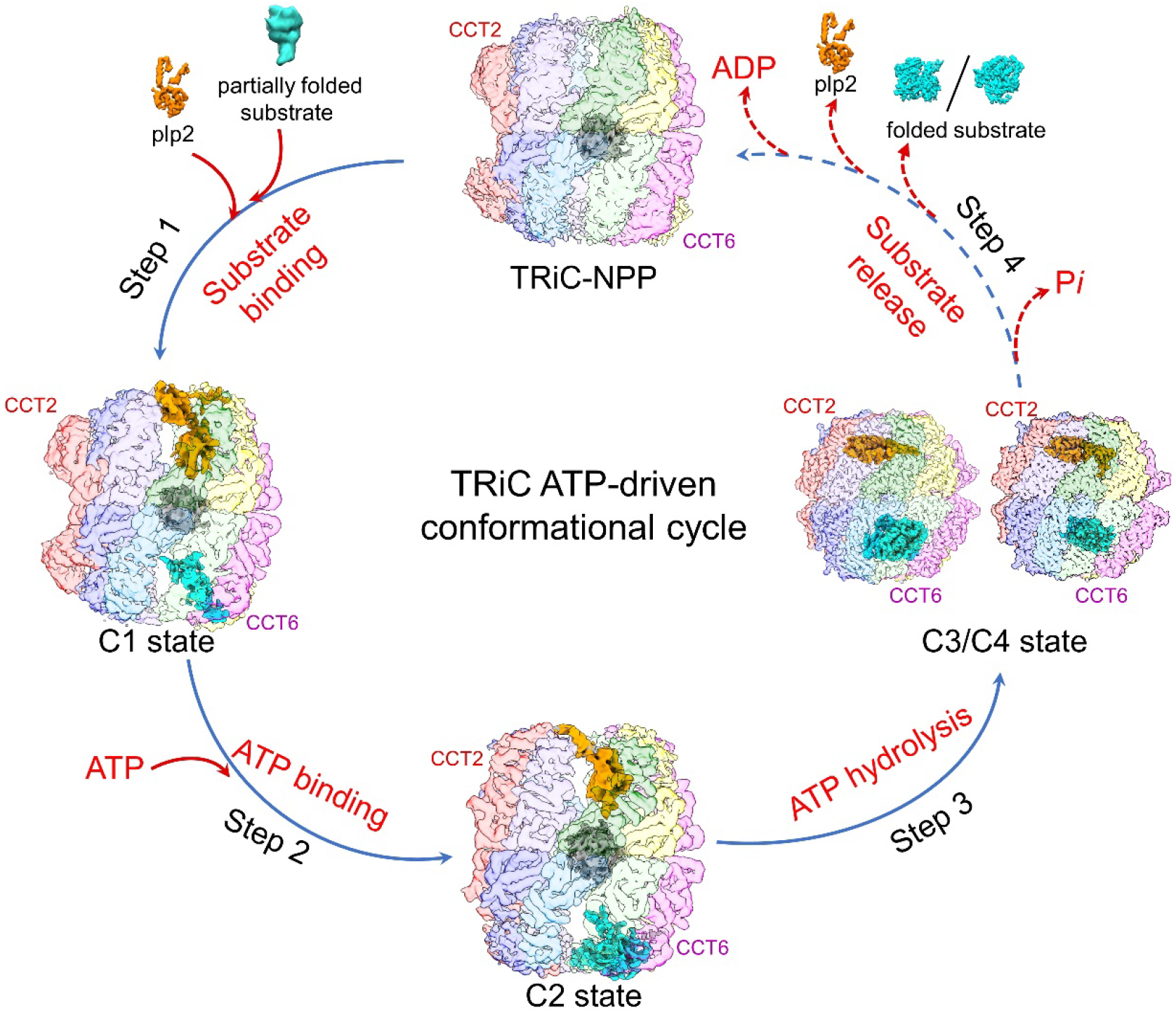
Proposed TRiC-plp2 mediated substrate folding process. Cochaperone plp2 and partially folded substrate can simultaneously bind open state TRiC within distinct chamber. Along with ATP binding, the location of substrate within TRiC begin to change, but plp2 remains in original location. Accompanying ATP hydrolysis and TRiC ring closure, both plp2 and substrate raise up to lie across the chamber, especially plp2, appearing tilting underneath the dome. Since the TRiC chamber reopen and the release of P*i*, plp2 and nearly folded substrate are released from the chamber, starting or waiting for a new round of substrate folding.

We found that in the open C1 (NPP) state, substrates tubulin/actin engage with the CCT6 hemisphere subunits (CCT8/6/3, Fig. 1D), and in the closed C3/C4 states, tubulin/actin remain in the CCT6 hemisphere (Fig. 3E and 4E). Consistently, previous studies also suggested other substrates (including AML1-ETO, Gβ, and mLST8) prefer to bind CCT6 hemisphere, especially in open-state TRiC (*51–53*). We have previously showed that in the NPP state TRiC, CCT8/6/3 in the CCT6 hemisphere have preloaded ADP from the environment of cells (*18*), which may stabilize these subunits making them serve as an anchor for substrate engagement. Collectively, the asymmetry or specificity of TRiC subunit in nucleotide binding may contribute to substrate engagement and processing.

### Potential roll of plp2 in TRiC-assisted tubulin/actin folding

Our data indicate that in yeast, plp2 coordinates with TRiC assisting the folding of major cytoskeletal proteins tubulin/actin, with plp2 serving as a cochaperone. Still, compared with other cochaperones, plp2 exhibits more complex interaction mode with TRiC in the following aspects: (1) plp2 engages with TRiC inside its chamber in both open and closed states (Fig. 6), while other cochaperones, such as Hsp70, PFD, and PhLP1, usually bind open state TRiC outside the chamber (*15, 28, 33, 44, 51, 54–56*); (2) plp2 and substrate occupy opposite TRiC chambers, instead of in the same ring as in the TRiC-prefoldin-actin and TRiC-PhLP1-Gβ cases (*44, 51*); (3) plp2 engages with TRiC through a combination of electrostatic and hydrophilic interactions (Fig. 5H-I), more complex than that of the electrostatic prefoldin-TRiC interaction (*44*). Intriguingly, along with TRiC ring closure, we captured plp2 translocation within TRiC chamber coordinating substrate translocation, not reported before, potentially facilitating substrate stabilization and folding.

Here, we captured the ternary complex of TRiC-plp2-tubulin (C3 state), with the full-length β-tubulin in the folded native state even loaded with a GTP (Fig. 3F). However, in the available TRiC-tubulin structures, plp2 is absent and tubulin is either with the I domain missing (*43*) or only partially ordered (*40*). This indicates the role of plp2 as a cochaperonin in tubulin stabilization and folding. Still, in our C4 map the actin density within TRiC was less well resolved compared with that of the closed human TRiC-actin-PhLP2A structure (*43*), in which actin was stabilized by the direct contacting long helix of the N-terminal domain of PhLP2A, not visualized in our C4 map. Nevertheless, the abovementioned differences could also be attributed to the distinct species, experimental conditions, or substrate folding stages, which need further examination.

We have previously shown that in yeast TRiC-ATP-AlFx free of substrate and cochaperone, almost all TRiC particles appear in the closed state (*19*). Here, our structural analysis demonstrated that the presence of plp2 and substrate could change the open-closed population ratio of TRiC, i.e. in such system 27% of TRiC particles remain open in the presence of ATP-AlFx (Fig. S3A-B), in line with a recent study on TRiC-prefoldin-σ3 (*28*). These results collectively indicate that the engagement of plp2 and substrate may slow down TRiC ring closure rate. Moreover, it has been proposed that being less cooperative than attainable allows chaperonins to support robust folding over a wider range of metabolic conditions (*57, 58*). Taken together, we postulate that with the engagement of plp2 and substrate, TRiC could tune its allosteric network to be less cooperative, slowing down its ring closure; this way substrate may have a higher chance of being folded to its native state inside TRiC chamber in a given round of cycling.

In summary, the present study reveals a more complete picture of plp2-mediated tubulin and actin folding assisted by TRiC accompanying its ATP-driven conformational cycle. Moreover, our study indicates that through engagement with TRiC via occupying one chamber throughout its ring closure process, plp2 may involve in mediating the substrate engagement and stabilization in the opposite TRiC chamber, and also in the substrate folding process through adjusting the ring closure rate of TRiC. Collectively, our data indicate that in yeast system, plp2 play a role as cochaperone in the TRiC-assisted folding of tubulin/actin. Our findings provide structural insights into the folding mechanism of the major cytoskeletal proteins tubulin/actin under the coordination of the complex biogenesis machinery TRiC and plp2, and could extend our understanding on the links between cytoskeletal proteostasis and human diseases such as developmental and neurological disorders.

## Acknowledgements

We are grateful to the staffs of the NCPSS Electron Microscopy facility, Database and Computing facility, Protein Expression and Purification facility, and Mass Spectrometry facility for instrument support and technical assistance. This work was supported by grants from the NSFC-ISF 31861143028, the Strategic Priority Research Program of CAS (XDB37040103), National Basic Research Program of China (2017YFA0503503), the NSFC (31670754 and 31872714 to Y.C, 31800623 to Z.D.), Shanghai Academic Research Leader (20XD1404200), and the CAS Facility-based Open Research Program and the CAS-Shanghai Science Research Center (CAS-SSRC-YH-2015-01, DSS-WXJZ-2018-0002).

## Author contributions

W.H., M.J. and Y.C. designed the experiments. W.H., Y.W. and M.J. purified the proteins, prepared the sample, and collected the cryo-EM data. W.H., YF.W. and C.L. performed data reconstruction. W.H. performed model building. W.H., Q.Z. and S.W. performed the Native Gel and ATPase experiments. C.P. and Y.Y. performed the XL-MS experiments and data analysis. W.H. and Y.C. analyzed the structure and wrote the manuscript.

## Competing interests

The authors declare no competing interests.

## Methods

### Purification of yeast TRiC

Yeast TRiC was purified according to our published protocol (*18*). The supernatant of yeast lysate was incubated with calmodulin resin (GE Healthcare) overnight at 4 °C. Elution of TRiC was achieved by using elution buffer (20 mM HEPES pH 7.5, 5 mM MgCl2, 0.1 mM EDTA, 50 mM NaCl, 1 mM DTT, 10% glycerol, 2 mM EGTA). The pooled eluate containing TRiC was further purified through size-exclusion chromatography (Superose 6 increase, GE Healthcare) and then concentrated with a Millipore Ultrafree centrifugal filter device (100-kDa cutoff).

### Expression and purification of yeast plp2 and actin

The yeast plp2 and actin were cloned into a modified pET28a vector that included a TEV protease site after the N-terminal 6xHis tag, and were expressed in *E. coli* Rosetta (DE3). Plp2 was purified by carrying out a one-step Ni affinity extraction, followed by size-exclusion chromatography (Superdex200, GE Healthcare). Actin was expressed as inclusion body. The pellet contains inclusion bodies were first denaturized by 6 M guanidine hydrochloride (GuHCl) and then renatured by dialyzing in dialysis buffer (25 mM Tris-HCl pH 7.5, 150 mM NaCl, 2 mM CaCl2, 0.25 mM ATP). The renaturing supernatant was purified through Ni affinity extraction, followed by size-exclusion chromatography (Superdex200, GE Healthcare).

### TRiC-plp2-actin complex formation

To prepare the sample of TRiC-plp2-actin without addition of nucleotide, we diluted the purified yeast TRiC with MQA buffer (20 mM HEPES, pH 7.5, 50 mM NaCl, 5 mM MgCl2, 0.1 mM EDTA, 1 mM DTT) to 1 μM, then incubated with plp2 and β-actin in the presence of 2 mM EDTA at 30 °C for 30 min, leading to a molar ratio of 1:10:10 among them. To prepare the sample of TRiC-plp2-actin in the presence of 1mM ATP-AlFx, we further incubated the abovementioned sample with 1 mM ATP, 10 mM MgCl2, 5 mM Al(NO3)3, and 30 mM NaF at 30 °C for 1hr. The same procedure was applied for the TRiC-actin with ATP-AlFx.

### Cross-linking and mass spectrometry analysis

The TRiC-plp2-actin complex was prepared as above, the complex was cross-linked by bis[sulfosuccinimidyl] suberate (BS3) (Sigma), with a spacer arm of 11.4 Å between their Cα carbons, on ice for 2 hours. The final concentration of crosslinker was at 2 mM. 50 mM Tris-HCl pH 7.5 was used to terminate the reaction at room temperature for 15 minutes. Cross-linked complexes were precipitated and digested for 16 hours at 37 °C by trypsin at an enzyme-to-substrate ratio of 1:50 (w/w). The tryptic digested peptides were desalted and loaded on an in-house packed capillary reverse-phase C18 column (40 cm length, 100 μM ID x 360 μM OD, 1.9 μM particle size, 120 Å pore diameter) connected to an Easy LC 1200 system. The samples were analyzed with a 120 min-HPLC gradient from 6% to 35% of buffer B (buffer A: 0.1% formic acid in Water; buffer B: 0.1% formic acid in 80% acetonitrile) at 300 nL/min. The eluted peptides were ionized and directly introduced into a Q-Exactive mass spectrometer using a nano-spray source. Survey full-scan MS spectra (from m/z 300-1800) were acquired in the Orbitrap analyzer with resolution r =70,000 at m/z 400. Cross-linked peptides were identified and evaluated using pLink2 software (*59*).

### Cryo-EM sample preparation and data acquisition

To prepare the vitrious sample of TRiC-plp2-actin without addition of nucleotide or TRiC-plp2-actin and TRiC-actin in the presence of ATP-AlFx, respectively, an aliquot of 2.2 μL biological sample obtained through abovementioned producer was applied to a glow-discharged holey carbon grid (R1.2/1.3, 200 mesh; Quantifoil). The grid was blotted with Vitrobot Mark IV (Thermo Fisher Scientific) and then plunged into liquid ethane cooled by liquid nitrogen. To handle the preferred orientation problem, the grid was pretreated with polylysine (*18*).

For each of the experimental sample conditions mentioned above, cryo-EM movies of the sample were collected on a Titan Krios electron microscope equipped with a Cs corrector (Thermo Fisher Scientific) operated at an accelerating voltage of 300 kV with a nominal magnification of 18,000x (Table S1). The movies were recorded on a K2 Summit direct electron detector (Gatan) operated in the super-resolution mode (yielding a pixel size of 1.318 Å after 2 times binning) under low-dose condition in an automatic manner using SerialEM (*60*). Each frame was exposed for 0.2 s and the total accumulation time was 7.6 s, leading to a total accumulated dose of 38 e^-^/Å^2^ on the specimen.

### Image processing and 3D reconstruction

The image processing and reconstruction procedure are shown in Fig. S2, S3, S6 and Table S1. For the TRiC-plp2-actin sample, the motion correction of each movie stack was performed using the Motioncorr2 program (*61*) before further image processing. The CTF parameters of each image were determined using CTFFIND4 (*62*). About 1000 particles were manually picked and then 2D classification using Relion 3.1, after which several good classes averages were chosen as template for further automatic particle picking in Relion3.1 (*63*). Bad particles and ice contaminations were then excluded by manual selection and 2D classification. Then 513,339 particles were refined and re-extract to re-center through a previous determined TRiC map, which was low-pass filtered to 20 Å. No symmetry was imposed in our procedure.

We first 3D classified the entire dataset into six classes: the last two classes presented better structural features (than the other four classes), and the remaining four classes were combined for another round of 3D classification, generating six classes, four of which showed better structural features and were combined for another round of 2D classification. Then the selected particles were combined with the previous good particles, and performed CTF refinement and per-particle motion correction (also known as Bayesian polishing) in Relion3.1. Then we masked extra densities out and subtracted particles for two rounds of focused 3D classification. 50,513 particles with better extra densities were selected for the final reconstruction to 4.55-Å map. The resolution estimation was based on the gold-standard Fourier Shell Correlation (FSC) 0.143 criterion, and the local resolution was estimated using Resmap (*25*).

For the cryo-EM samples of TRiC-plp2-actin with ATP-AlFx, similar procedures were adopted as above. For more details, please refer to supplementary figure S3, and Table S1. Moreover, for the open dataset, after obtained a 3.78-Å map, we also applied focused 3D classification on the extra density of cis-/trans-ring, respectively. 33,866 particles were reconstructed to a final 3.91-Å map with extra densities in both rings. For the closed dataset, after obtained a 2.97-Å map using the above-mentioned procedure, we masked extra densities out and subtracted particles for focused 3D classification. Two new classes had different densities in both rings with 10.30% and 12.38% particles, respectively. While the major particles with 69.67% particles had the same conformation in both rings. Then we reverted the two new classes to original particles for refinement. For exhaustive classification of the extra density, we subtracted cis-ring and trans-ring of closed TRiC particles, respectively. We then applied focused 3D classification with different mask on cis-ring or trans-ring particles. Through combining the classified particles of cis-ring and trans-ring and taking the intersection of these classes (different colored box line represented the same selection of the intersection), three new conformations with distinct extra densities in trans-ring were resolved at 3.39-, 3.20- and 3.05-Å, termed as “C3”, “C4” and “C5” state, respectively. The resolution estimation was based on the gold-standard Fourier Shell Correlation (FSC) 0.143 criterion, and the local resolution was estimated using Resmap (*25*).

For the TRiC-actin dataset, please refer to Fig. S6, 94,912 particles were remained after 2D classification. After several rounds of 2D/3D classification, 17,156 particles at closed state of TRiC were selected and loaded into cryoSPARC V3.3.1, final particles were refined to 3.87-Å-resolution, using Non-uniform refinement. The overall resolution was determined based on the gold-standard criterion using a Fourier shell correlation (FSC) of 0.143.

### Pseudo atomic model building by flexible fitting

We built pseudo atomic model for the better resolved C1, C2, C3, C4 and C5 maps. TRiC-NPP (PDB: 5GW4), TRiC-AMP-PNP (PDB: 5GW5) and closed TRiC (PDB: 6KS6), which were more conformationally similar to our current structures, were used as the initial model. Then the models for the subunits were fitted into the density map simultaneously as a rigid body using UCSF Chimera and were combined as a complete model (*64*). Later, we used *phenix.real_space_refine* to improve the fitting (*65*).

Furthermore, we performed real space refinement using COOT (*66*) in order to eliminate steric clashes, and the last round of flexible fitting on the entire complex using Rosetta followed by Phenix (*67*).

For the plp2 model in C3, C4 and C5, we first chose the complete model of yeast plp2 from database of AlphaFold (*68*). After rigid body docking, we performed flexible fitting on each of the domain using *phenix.real_space_refine* and one more round of flexible fitting using Rosetta. Finally, we performed real space refinement using COOT in order to eliminate steric clashes, and the last round of flexible fitting of plp2 using Rosetta followed by Phenix.

For the tubulin model in C3, we first chose the complete model of yeast β-tubulin (PDB:5W3F) as initial model. After rigid body docking, we performed flexible fitting on each of the domain using *phenix.real_space_refine* and one more round of flexible fitting using Rosetta. Finally, we performed real space refinement using COOT in order to eliminate steric clashes, and the last round of flexible fitting of tubulin using Rosetta followed by Phenix.

### ATPase activity assay

ATP hydrolysis rates of TRiC with plp2 and β-actin were measured by performing an NADH coupled assay (*69*). In this assay, each ATP hydrolysis event allows conversion of one molecule of phosphoenolpyruvate into pyruvate by pyruvate kinase. Thereafter, pyruvate is converted to lactate by L-lactate dehydrogenase, which results in oxidation of a single NADH molecule. Loss of NADH over time, which is quantifiably proportional to ATP hydrolysis rates, is monitored by measuring the decrease in absorbance at 340 nm. All of the assays were conducted at 30°C in a buffer containing 10 mM Mops-KOH, pH 8.0, 20 mM KCl, and 10 mM MgCl2, and in the presence of 1 mM ATP. Experiments were performed in triplicate using 1 μM of the protein complex. Absorbance was measured in a 100 μl reaction volume using a 96-well plate reader. Data analysis was performed using Graphpad Prism 8.

### Thermal stability measurement

To test the thermal stability of plp2 and β-actin, proteins alone or with 2mM EDTA pretreated were applied into capillary tube, and then put them into Prometheus (Nano Temper) for thermal stability measurement. Meanwhile, we used α-actin from rabbit muscle (Sigma) for control. The temperature was increased 2.0°C per minutes until 95°C. Then the thermal stability curves were described through the temperature on X-axis and first derivative of the ratio between integrated fluorescence at 350nm and 330nm on Y-axis. Each experiment was repeated for three times.

## Supplemental figures

**Fig. S1.**
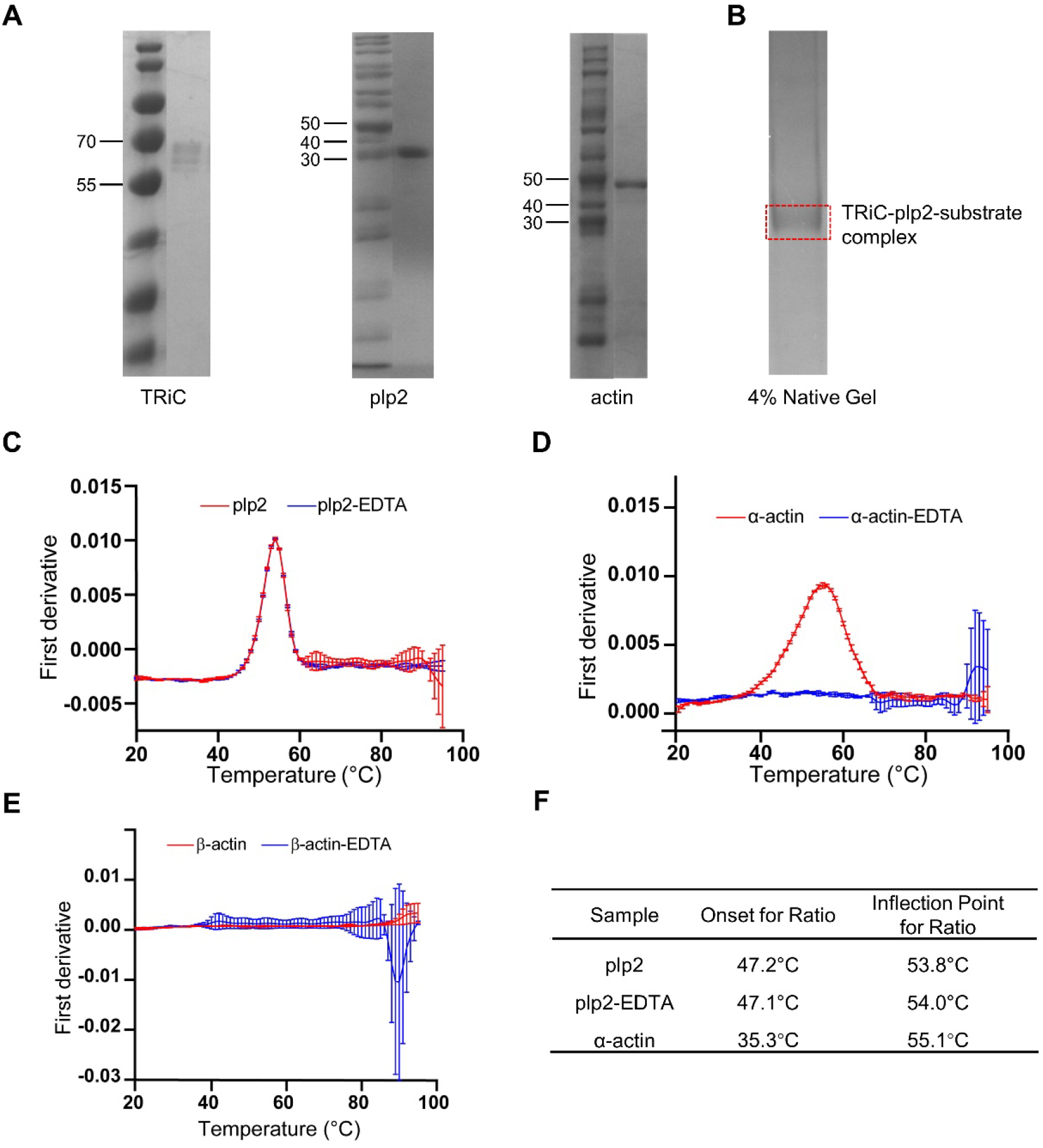
Protein purification and identification. (A) Protein purification of yeast TRiC, yeast plp2 and yeast β-actin from left to right. (B) Association of TRiC-plp2-actin monitored by 4% native gel, the complex was indicated in red box which identified by mass spectrum (Table S2). (C-E) Measurements of the thermal stability among plp2 (C), α-actin from rabbit muscle (D) and yeast β-actin (E). (F) Table of the thermal stability results showed the first derivative of the ratio between integrated fluorescence at 350nm and 330nm on Y-axis.

**Fig. S2.**
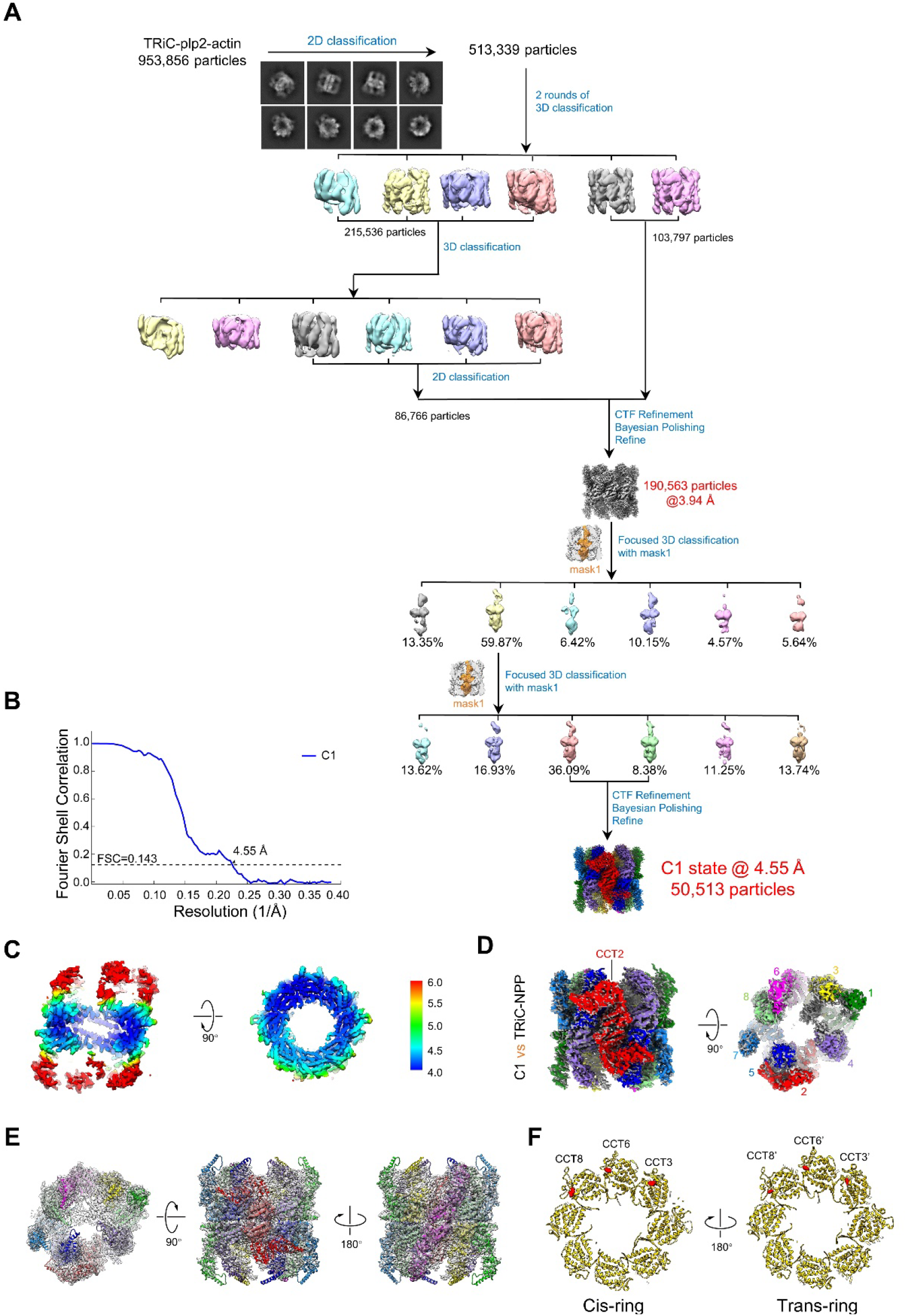
Data processing of TRiC-plp2-substrate. (A) cryo-EM data processing procedure for the TRiC-plp2-substrate dataset. (B) Resolution estimations of C1 map according to the gold-standard FSC criterion of 0.143. (C) Local resolution evaluation for the C1 map. (D) Comparison between C1 map (grey) and TRiC-NPP (EMD-9540). (E) Model map fitting of C1 map. (F) Nucleotide state of C1 map.

**Fig. S3.**
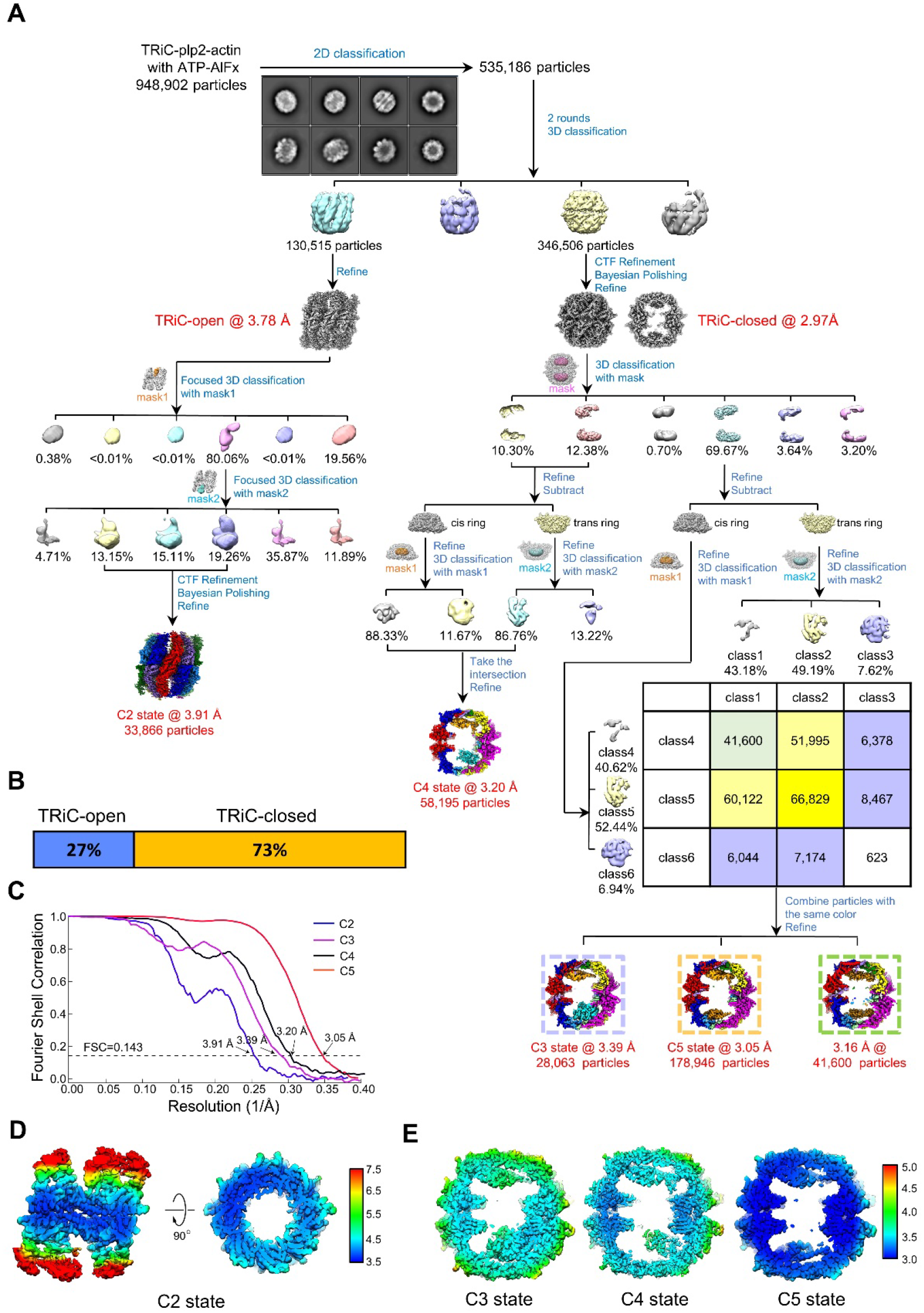
Data processing of TRiC-plp2-substrate with ATP-AlFx. (A) cryo-EM data processing procedure for the TRiC-plp2-substrate with ATP-AlFx. (B) Population distribution of TRiC-plp2-substrate with ATP-AlFx. (C) Resolution estimations of C2/3/4/5 maps according to the gold-standard FSC criterion of 0.143. (D-E) Local resolution estimations for the C2/3/4/5 maps.

**Fig. S4.**
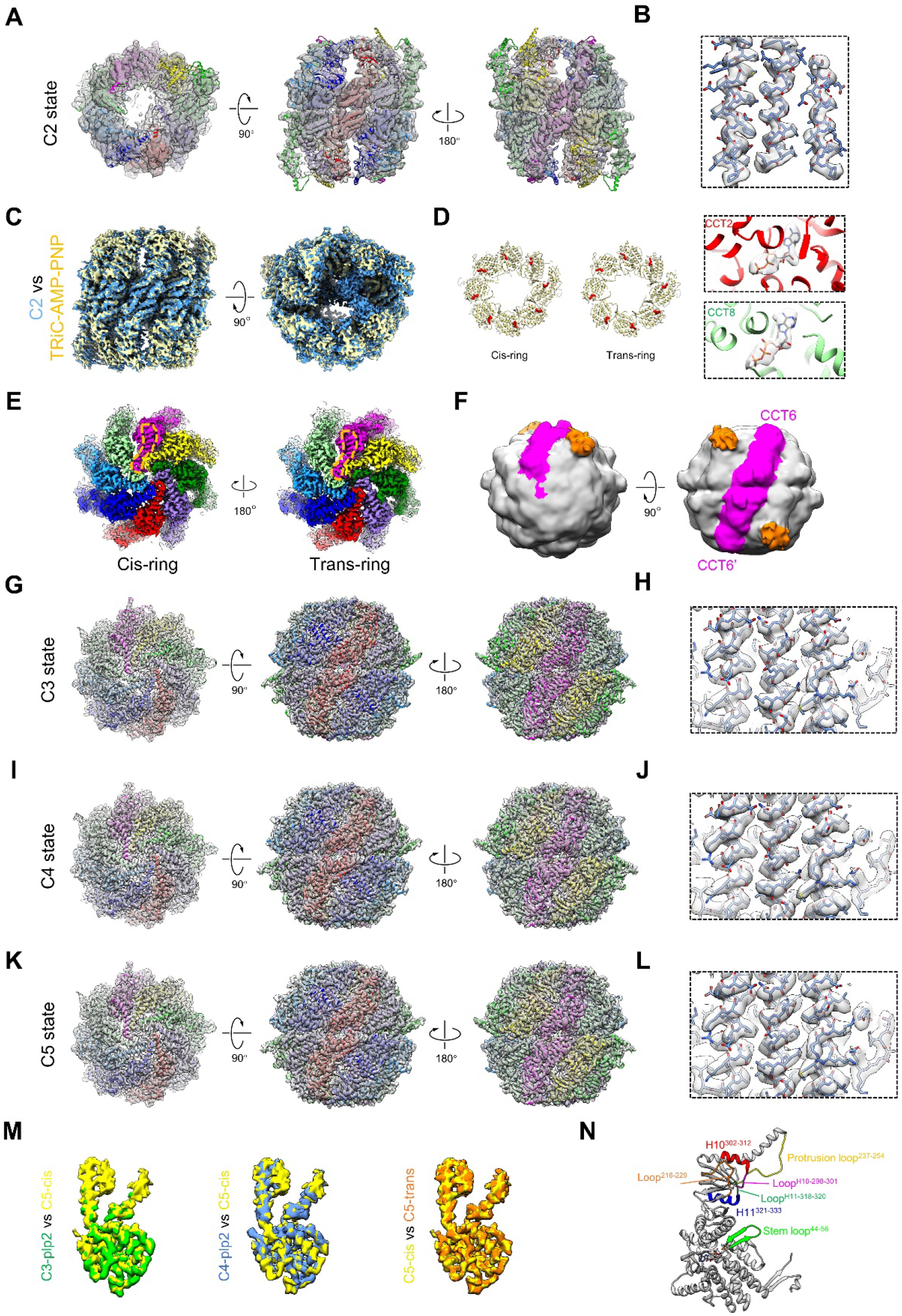
Pseudoatomic model of TRiC-plp2-substrate with ATP-AlFx. (A) Model and map fitting of C2 state. (B) Representative atomic-resolution structural features of C2 map. (C) Comparison between C2 map and TRiC-AMP-PNP (EMD-9541). (D) Nucleotide occupancy statuses of C2 map, and representative nucleotide density (in the ATP binding state) of CCT2 and CCT8 in C2 map. (E) The unique kink feature (enclosed in orange dashed line) in the apical domain of Helix 9 of CCT6 were unambiguously visualized in both rings of closed TRiC (C5 map), substantiating the on-axis location of CCT6. (F) Direct visualization of the protruding CBP tag (orange, fused to the A domain of CCT6) on closed TRiC. (G-L) Model-map fitting and representative atomic-resolution structural features of C3/C4/C5 map. (M) Plp2 densities comparison among C3, C4 and C5 showed they all exhibited the same conformation and location. (N) Key structural elements of TRiC subunit involved in the interaction with tubulin, with the residue numbers of these elements being labeled (CCT6 as a representative subunit).

**Fig. S5.**
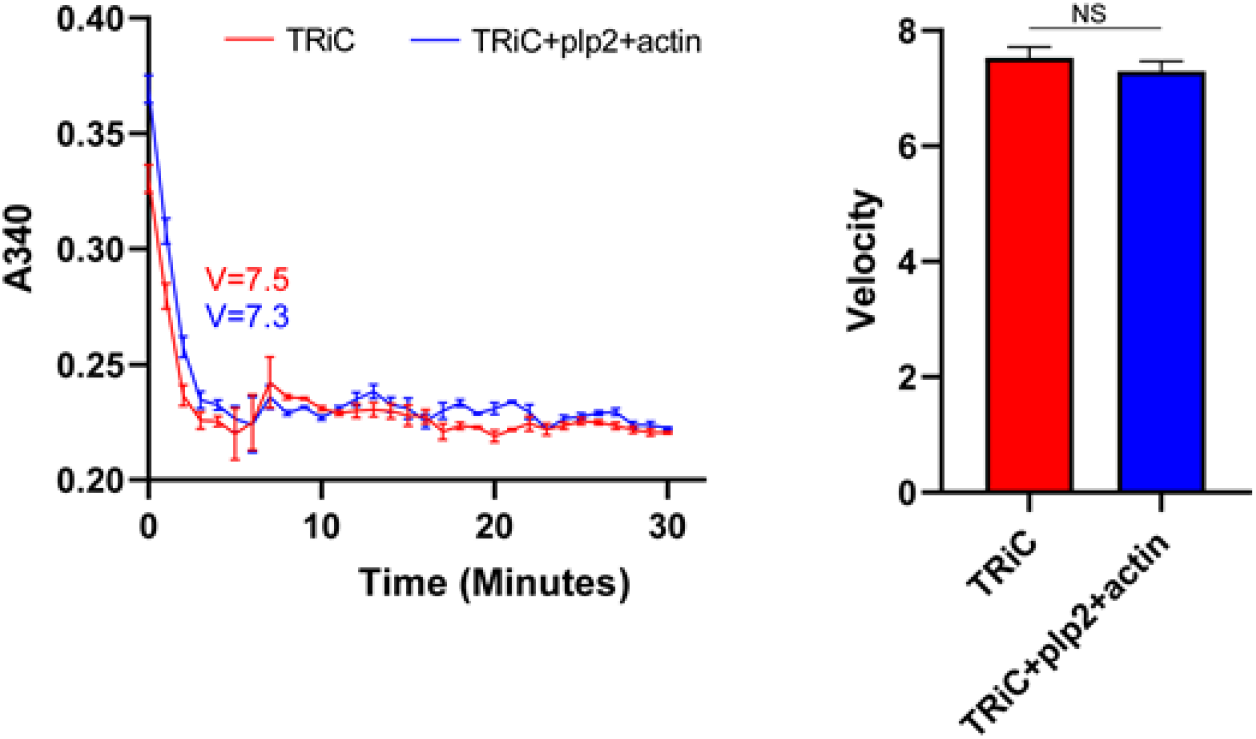
NADH-coupled assay identified that the existence of plp2 and actin had no significant effect on ATPase activity of TRiC. We chose first three point to calculate the velocity, and each experiment was repeated three times.

**Fig. S6.**
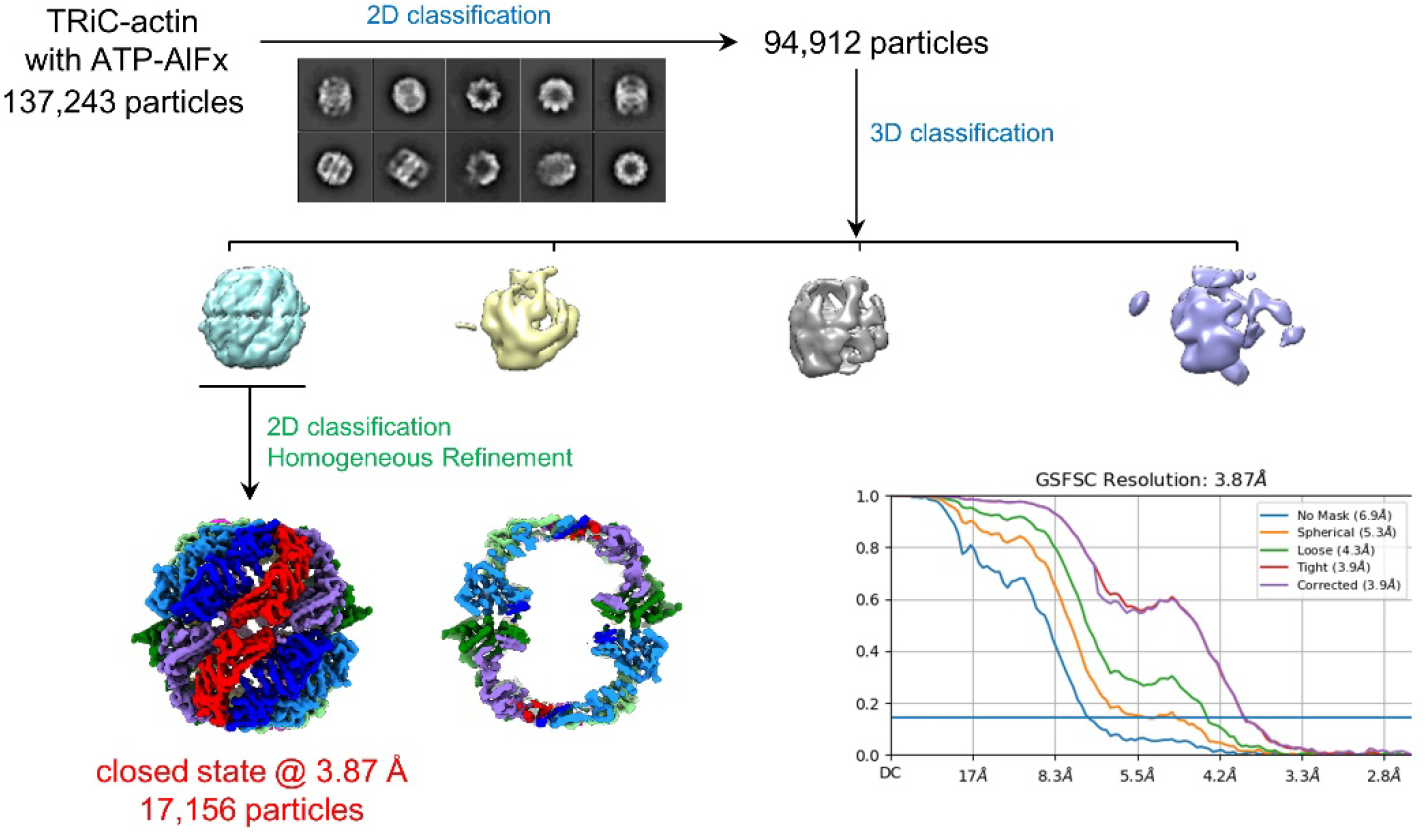
Cryo-EM data processing procedure for TRiC-actin with ATP-AlFx. Data processing of TRiC-actin with ATP-AlFx found no extra densities in closed TRiC chamber without the presence of TRiC. FSC curve at 0.143 criterion shows an overall resolution of closed map.

## Supplemental tables

**Table S1.**
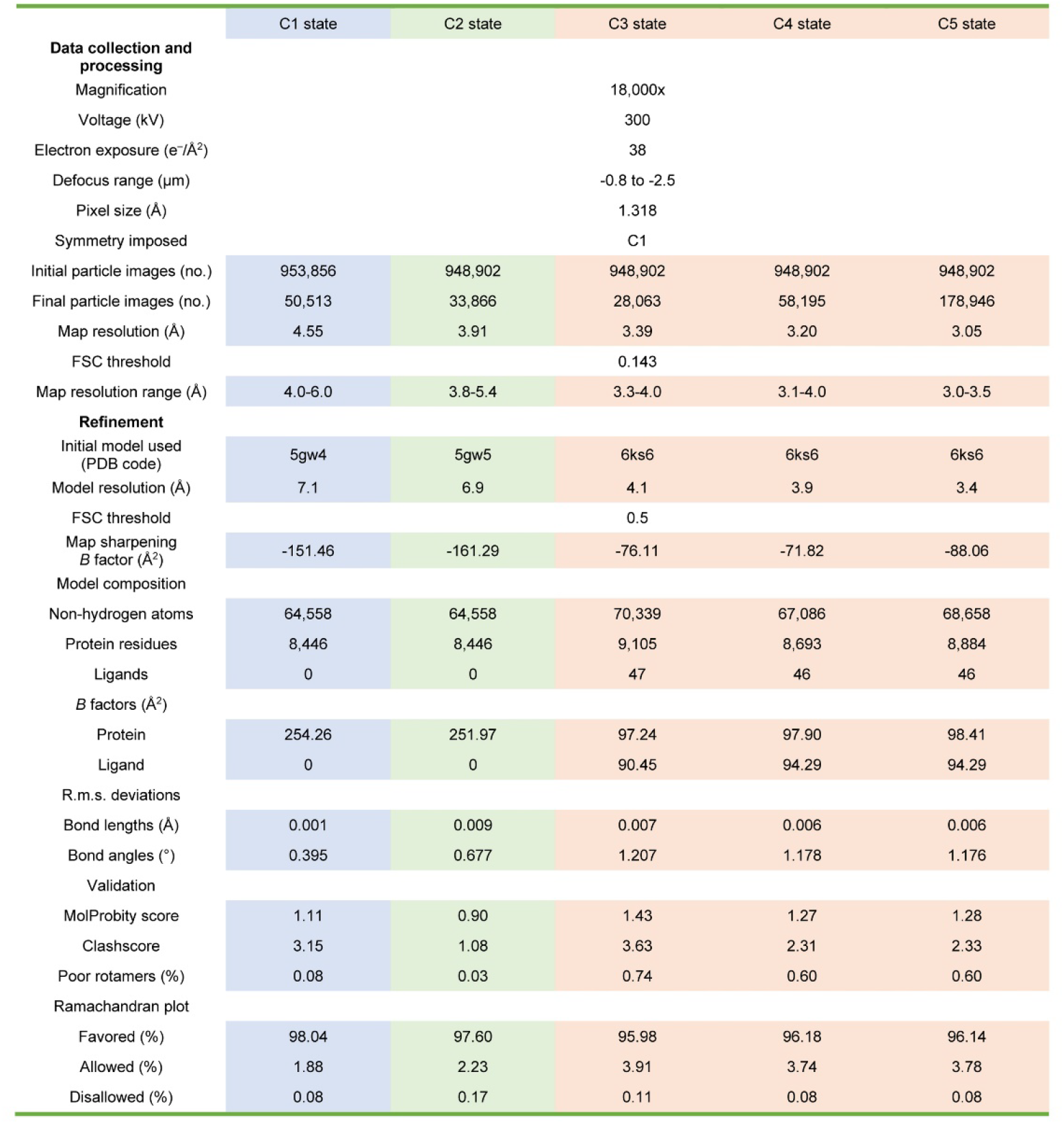
Statistics of cryo-EM data collection, processing, and model validation.

|  | C1 state | C2 state | C3 state | C4 state | C5 state |
| --- | --- | --- | --- | --- | --- |
| **Data collection and processing** |  |  |  |  |  |
| Magnification |  |  | 18,000x |  |  |
| Voltage (kV) |  |  | 300 |  |  |
| Electron exposure (e<sup>-</sup>/Å<sup>2</sup>) |  |  | 38 |  |  |
| Defocus range (μm) |  |  | -0.8 to -2.5 |  |  |
| Pixel size (Å) |  |  | 1.318 |  |  |
| Symmetry imposed |  |  | C1 |  |  |
| Initial particle images (no.) | 953,856 | 948,902 | 948,902 | 948,902 | 948,902 |
| Final particle images (no.) | 50,513 | 33,866 | 28,063 | 58,195 | 178,946 |
| Map resolution (Å) | 4.55 | 3.91 | 3.39 | 3.20 | 3.05 |
| FSC threshold |  |  | 0.143 |  |  |
| Map resolution range (Å) | 4.0-6.0 | 3.8-5.4 | 3.3-4.0 | 3.1-4.0 | 3.0-3.5 |
| **Refinement** |  |  |  |  |  |
| Initial model used (PDB code) | 5gw4 | 5gw5 | 6ks6 | 6ks6 | 6ks6 |
| Model resolution (Å) | 7.1 | 6.9 | 4.1 | 3.9 | 3.4 |
| FSC threshold |  |  | 0.5 |  |  |
| Map sharpening B factor (Å<sup>2</sup>) | -151.46 | -161.29 | -76.11 | -71.82 | -88.06 |
| **Model composition** |  |  |  |  |  |
| Non-hydrogen atoms | 64,558 | 64,558 | 70,339 | 67,086 | 68,658 |
| Protein residues | 8,446 | 8,446 | 9,105 | 8,693 | 8,884 |
| Ligands | 0 | 0 | 47 | 46 | 46 |
| **B factors (Å<sup>2</sup>)** |  |  |  |  |  |
| Protein | 254.26 | 251.97 | 97.24 | 97.90 | 98.41 |
| Ligand | 0 | 0 | 90.45 | 94.29 | 94.29 |
| **R.m.s. deviations** |  |  |  |  |  |
| Bond lengths (Å) | 0.001 | 0.009 | 0.007 | 0.006 | 0.006 |
| Bond angles (°) | 0.395 | 0.677 | 1.207 | 1.178 | 1.176 |
| **Validation** |  |  |  |  |  |
| MolProbity score | 1.11 | 0.90 | 1.43 | 1.27 | 1.28 |
| Clashscore | 3.15 | 1.08 | 3.63 | 2.31 | 2.33 |
| Poor rotamers (%) | 0.08 | 0.03 | 0.74 | 0.60 | 0.60 |
| **Ramachandran plot** |  |  |  |  |  |
| Favored (%) | 98.04 | 97.60 | 95.98 | 96.18 | 96.14 |
| Allowed (%) | 1.88 | 2.23 | 3.91 | 3.74 | 3.78 |
| Disallowed (%) | 0.08 | 0.17 | 0.11 | 0.08 | 0.08 |

**Table S2.** MS of native gel identified TRiC cooperated with plp2, actin and tubulin.

|  | Protein | Coverage | PSMs | Peptides | Unique peptides |
| --- | --- | --- | --- | --- | --- |
| TRiC subunits | CCT1 | 93 | 408 | 64 | 64 |
|  | CCT2 | 89 | 463 | 61 | 61 |
|  | CCT3 | 84 | 388 | 62 | 62 |
|  | CCT4 | 81 | 387 | 55 | 55 |
|  | CCT5 | 82 | 364 | 62 | 62 |
|  | CCT6 | 83 | 385 | 58 | 58 |
|  | CCT7 | 91 | 369 | 59 | 59 |
|  | CCT8 | 88 | 526 | 65 | 65 |
| Other proteins | plp2 | 81 | 200 | 29 | 29 |
|  | actin | 74 | 72 | 25 | 25 |
| | β-tubulin | 41.35 | 18 | 14 | 14 |

**Table S3.** XL-MS analysis of TRiC with plp2 and actin.

| protein1-protein2 | peptides | E-value | Spec count |
| --- | --- | --- | --- |
| CCT1(529)-plp2(26) | SLKSALEACVAILR(3)-<br>AKGVIPER(2) | 1.88E-04 | 2 |
| CCT2(270)-plp2(104) | EKMKNK(4)-<br>FGEV FHINKPEYNK(9) | 2.45E-02 | 28 |
| CCT3(126)-plp2(248) | NIHPV IIIQALKK(12)-<br>KLHYGEK(1) | 1.77E-23 | 13 |
| CCT3(253)-plp2(95) | VLLDCPLEYKK(11)-<br>SKFGEV FHINKPEYNK(2) | 2.39E-02 | 6 |
| CCT4(276)-plp2(26) | AYLLNICKK(8)-<br>AKGVIPER(2) | 1.02E-10 | 2 |
| CCT8(322)-plp2(26) | YGILVLKVPSK(7)-<br>AKGVIPER(2) | 1.19E-02 | 2 |
| actin(315)-plp2(26) | MQKEITALAPSSMK(3)-<br>AKGVIPER(2) | 1.50E-03 | 4 |
| actin(328)-plp2(26) | VKIIAPPER(2)-<br>AKGVIPER(2) | 2.61E-05 | 2 |
| actin(315)-CCT6(40) | MQKEITALAPSSMK(3)-<br>VNV TSAEGLQSVLETNLG<br>PKGTLK(20) | 5.38E-02 | 3 |

\* We used E-value (1.00E-02) and spec count (larger than 1) as the threshold to remove extra XL-MS data with lower confidence.

**Table S4.**
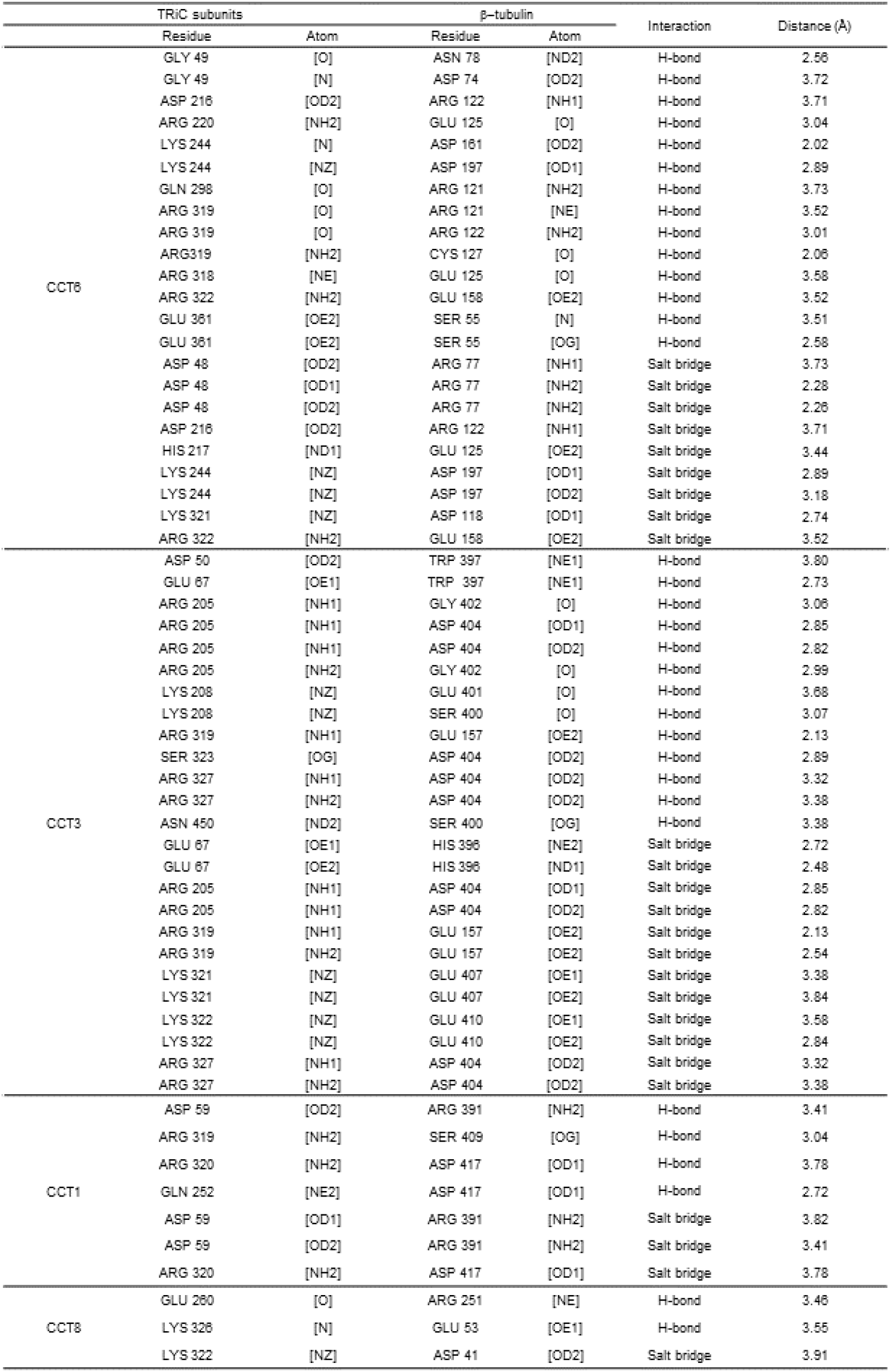
Interactions between closed TRiC and tubulin analyzed by PISA.

**Table S5.** Interactions between closed TRiC and plp2 analyzed by PISA.

|  | TRiC subunits |  | plp2 |  | Interaction | Distance (Å) |
| --- | --- | --- | --- | --- | --- | --- |
|  | Residue | Atom | Residue | Atom |  |  |
| CCT5 | GLU 277 | [OE2] | ASN 103 | [ND2] | H-bond | 2.13 |
|  | GLN 324 | [O] | LYS 104 | [NZ] | H-bond | 2.74 |
|  | LYS 247 | [NZ] | GLU 179 | [OE1] | Salt bridge | 2.53 |
|  | GLU 277 | [OE1] | ARG 176 | [NH1] | Salt bridge | 3.71 |
|  | LYS 282 | [NZ] | GLU 106 | [OE1] | Salt bridge | 2.47 |
|  | LYS 282 | [NZ] | GLU 106 | [OE2] | Salt bridge | 2.48 |
|  | LYS 284 | [NZ] | GLU 98 | [OE1] | Salt bridge | 2.87 |
|  | LYS 286 | [NZ] | GLU 98 | [OE1] | Salt bridge | 3.38 |
| CCT2 | GLN 291 | [OH] | GLU 145 | [O] | H-bond | 3.30 |
|  | TYR 291 | [OH] | ARG 151 | [NH1] | H-bond | 2.42 |
|  | TYR 287 | [NH2] | GLN 183 | [OE2] | Salt bridge | 2.74 |
| CCT6 | TYR 242 | [OH] | GLN 91 | [OE1] | H-bond | 3.02 |
|  | VAL 247 | [O] | ARG 165 | [NH1] | H-bond | 3.83 |
|  | ASN 248 | [O] | ARG 165 | [NH1] | H-bond | 3.83 |
|  | ARG 322 | [N] | ARG 73 | [OE2] | H-bond | 2.24 |
|  | ARG 323 | [N] | GLU 73 | [OE1] | H-bond | 3.35 |
|  | LYS 321 | [NZ] | ASP 74 | [OD2] | Salt bridge | 3.69 |
|  | ARG 322 | [NH2] | GLU 68 | [OE2] | Salt bridge | 3.39 |
| CCT7 | LEU 250 | [O] | LYS 109 | [NZ] | H-bond | 3.76 |
|  | GLU 249 | [OE2] | LYS 109 | [NZ] | H-bond | 3.09 |
| CCT1 | ARG 319 | [NH1] | MET 246 | [OE2] | Salt bridge | 2.45 |
|  | ARG 319 | [NH2] | MET 246 | [OE1] | Salt bridge | 3.10 |
|  | ARG 320 | [NH1] | GLU 242 | [OE1] | Salt bridge | 3.21 |
|  | ARG 320 | [NE] | GLU 242 | [OE2] | Salt bridge | 2.44 |
| CCT3 | ARG 318 | [NE] | ASP 69 | [OD2] | Salt bridge | 3.38 |
|  | ARG 319 | [NH1] | ASP 69 | [OD1] | Salt bridge | 3.71 |
|  | ARG 319 | [NH2] | GLU 65 | [OE1] | Salt bridge | 3.52 |
|  | LYS 322 | [NZ] | GLU 245 | [OE2] | Salt bridge | 2.10 |

## Notes

### Competing Interest Statement

The authors have declared no competing interest.

## References

1. L. Chen et al., Chaperonin CCT-mediated AIB1 folding promotes the growth of ERalpha-positive breast cancer cells on hard substrates. PLoS One 9, e96085 (2014).

2. R. Bassiouni et al., Chaperonin Containing TCP-1 Protein Level in Breast Cancer Cells Predicts Therapeutic Application of a Cytotoxic Peptide. Clin Cancer Res 22, 4366–4379 (2016).

3. M. Jin, C. Liu, W. Han, Y. Cong, TRiC/CCT Chaperonin: Structure and Function. Subcell Biochem 93, 625–654 (2019).

4. R. Melki, N. J. Cowan, Facilitated folding of actins and tubulins occurs via a nucleotide-dependent interaction between cytoplasmic chaperonin and distinctive folding intermediates. Mol Cell Biol 14, 2895–2904 (1994).

5. D. B. Vinh, D. G. Drubin, A yeast TCP-1-like protein is required for actin function in vivo. Proceedings of the National Academy of Sciences of the United States of America 91, 9116–9120 (1994).

6. O. Llorca et al., Eukaryotic chaperonin CCT stabilizes actin and tubulin folding intermediates in open quasi-native conformations. The EMBO journal 19, 5971–5979 (2000).

7. O. Llorca et al., The ‘sequential allosteric ring’ mechanism in the eukaryotic chaperonin-assisted folding of actin and tubulin. The EMBO journal 20, 4065–4075 (2001).

8. A. Camasses, A. Bogdanova, A. Shevchenko, W. Zachariae, The CCT Chaperonin Promotes Activation of the Anaphase-Promoting Complex through the Generation of Functional Cdc20. Molecular Cell 12, 87–100 (2003).

9. A. G. Trinidad et al., Interaction of p53 with the CCT complex promotes protein folding and wild-type p53 activity. Mol Cell 50, 805–817 (2013).

10. M. Kasembeli et al., Modulation of STAT3 folding and function by TRiC/CCT chaperonin. PLoS Biol 12, e1001844 (2014).

11. A. J. McClellan, M. D. Scott, J. Frydman, Folding and quality control of the VHL tumor suppressor proceed through distinct chaperone pathways. Cell 121, 739–748 (2005).

12. Y. Cong et al., Symmetry-free cryo-EM structures of the chaperonin TRiC along its ATPase-driven conformational cycle. The EMBO journal 31, 720–730 (2012).

13. A. S. Meyer et al., Closing the folding chamber of the eukaryotic chaperonin requires the transition state of ATP hydrolysis. Cell 113, 369–381 (2003).

14. C. Spiess, A. S. Meyer, S. Reissmann, J. Frydman, Mechanism of the eukaryotic chaperonin: protein folding in the chamber of secrets. Trends Cell Biol 14, 598–604 (2004).

15. M. Jin et al., An ensemble of cryo-EM structures of TRiC reveal its conformational landscape and subunit specificity. Proceedings of the National Academy of Sciences of the United States of America 116, 19513–19522 (2019).

16. N. Kalisman, C. M. Adams, M. Levitt, Subunit order of eukaryotic TRiC/CCT chaperonin by cross-linking, mass spectrometry, and combinatorial homology modeling. Proceedings of the National Academy of Sciences of the United States of America 109, 2884–2889 (2012).

17. H. Wang, W. Han, J. Takagi, Y. Cong, Yeast Inner-Subunit PA–NZ-1 Labeling Strategy for Accurate Subunit Identification in a Macromolecular Complex through Cryo-EM Analysis. Journal of Molecular Biology 430, 1417–1425 (2018).

18. Y. Zang et al., Staggered ATP binding mechanism of eukaryotic chaperonin TRiC (CCT) revealed through high-resolution cryo-EM. Nat Struct Mol Biol 23, 1083–1091 (2016).

19. Y. Zang et al., Development of a yeast internal-subunit eGFP labeling strategy and its application in subunit identification in eukaryotic group II chaperonin TRiC/CCT. Scientific reports 8, 2374 (2018).

20. A. Leitner et al., The molecular architecture of the eukaryotic chaperonin TRiC/CCT. Structure 20, 814–825 (2012).

21. S. Reissmann et al., A gradient of ATP affinities generates an asymmetric power stroke driving the chaperonin TRIC/CCT folding cycle. Cell reports 2, 866–877 (2012).

22. L. Ditzel et al., Crystal structure of the thermosome, the archaeal chaperonin and homolog of CCT. Cell 93, 125–138 (1998).

23. T. Waldmann, A. Lupas, J. Kellermann, J. Peters, W. Baumeister, Primary structure of the thermosome from Thermoplasma acidophilum. Biol Chem Hoppe Seyler 376, 119–126 (1995).

24. M. Klumpp, W. Baumeister, L. O. Essen, Structure of the substrate binding domain of the thermosome, an archaeal group II chaperonin. Cell 91, 263–270 (1997).

25. V. F. Lundin, M. R. Leroux, P. C. Stirling, Quality control of cytoskeletal proteins and human disease. Trends Biochem Sci 35, 288–297 (2010).

26. I. E. Vainberg et al., Prefoldin, a chaperone that delivers unfolded proteins to cytosolic chaperonin. Cell 93, 863–873 (1998).

27. S. Geissler, K. Siegers, E. Schiebel, A novel protein complex promoting formation of functional alpha- and gamma-tubulin. The EMBO journal 17, 952–966 (1998).

28. J. J. Knowlton et al., Structural and functional dissection of reovirus capsid folding and assembly by the prefoldin-TRiC/CCT chaperone network. Proceedings of the National Academy of Sciences 118, (2021).

29. D. Gestaut et al., Structural visualization of the tubulin folding pathway directed by eukaryotic chaperonin TRiC. bioRxiv, (2022).

30. K. Siegers et al., Compartmentation of protein folding in vivo: sequestration of non-native polypeptide by the chaperonin-GimC system. The EMBO journal 18, 75–84 (1999).

31. R. Siegert, M. R. Leroux, C. Scheufler, F. U. Hartl, I. Moarefi, Structure of the molecular chaperone prefoldin: unique interaction of multiple coiled coil tentacles with unfolded proteins. Cell 103, 621–632 (2000).

32. B. M. Willardson, A. C. Howlett, Function of phosducin-like proteins in G protein signaling and chaperone-assisted protein folding. Cell Signal 19, 2417–2427 (2007).

33. P. C. Stirling et al., PhLP3 modulates CCT-mediated actin and tubulin folding via ternary complexes with substrates. The Journal of biological chemistry 281, 7012–7021 (2006).

34. C. Bregier et al., PHLP2 is essential and plays a role in ciliogenesis and microtubule assembly in Tetrahymena thermophila. J Cell Physiol 228, 2175–2189 (2013).

35. P. C. Stirling et al., Functional interaction between phosducin-like protein 2 and cytosolic chaperonin is essential for cytoskeletal protein function and cell cycle progression. Molecular biology of the cell 18, 2336–2345 (2007).

36. E. A. McCormack, G. M. Altschuler, C. Dekker, H. Filmore, K. R. Willison, Yeast phosducin-like protein 2 acts as a stimulatory co-factor for the folding of actin by the chaperonin CCT via a ternary complex. Journal of molecular biology 391, 192–206 (2009).

37. S. F. Stuart, R. J. Leatherbarrow, K. R. Willison, A two-step mechanism for the folding of actin by the yeast cytosolic chaperonin. The Journal of biological chemistry 286, 178–184 (2011).

38. K. R. Willison, The structure and evolution of eukaryotic chaperonin-containing TCP-1 and its mechanism that folds actin into a protein spring. Biochem J 475, 3009–3034 (2018).

39. D. Balchin, G. Milicic, M. Strauss, M. Hayer-Hartl, F. U. Hartl, Pathway of Actin Folding Directed by the Eukaryotic Chaperonin TRiC. Cell 174, 1507–1521 e1516 (2018).

40. I. G. Munoz et al., Crystal structure of the open conformation of the mammalian chaperonin CCT in complex with tubulin. Nat Struct Mol Biol 18, 14–19 (2011).

41. C. Dekker et al., The crystal structure of yeast CCT reveals intrinsic asymmetry of eukaryotic cytosolic chaperonins. The EMBO journal 30, 3078–3090 (2011).

42. O. Llorca et al., Eukaryotic type II chaperonin CCT interacts with actin through specific subunits. Nature 402, 693–696 (1999).

43. J. J. Kelly et al., Snapshots of actin and tubulin folding inside the TRiC chaperonin. Nat Struct Mol Biol (2022).

44. D. Gestaut et al., The Chaperonin TRiC/CCT Associates with Prefoldin through a Conserved Electrostatic Interface Essential for Cellular Proteostasis. Cell 177, 751–765 e715 (2019).

45. C. R. Booth et al., Mechanism of lid closure in the eukaryotic chaperonin TRiC/CCT. Nat Struct Mol Biol 15, 746–753 (2008).

46. Y. Cong et al., 4.0-A resolution cryo-EM structure of the mammalian chaperonin TRiC/CCT reveals its unique subunit arrangement. Proceedings of the National Academy of Sciences of the United States of America 107, 4967–4972 (2010).

47. E. Nogales, S. G. Wolf, K. H. Downing, Structure of the alpha beta tubulin dimer by electron crystallography. Nature 391, 199–203 (1998).

48. J. Lowe, H. Li, K. H. Downing, E. Nogales, Refined structure of alpha beta-tubulin at 3.5 A resolution. Journal of molecular biology 313, 1045–1057 (2001).

49. K. I. Brackley, J. Grantham, Activities of the chaperonin containing TCP-1 (CCT): implications for cell cycle progression and cytoskeletal organisation. Cell Stress Chaperones 14, 23–31 (2009).

50. G. W. Farr, E. C. Scharl, R. J. Schumacher, S. Sondek, A. L. Horwich, Chaperonin-mediated folding in the eukaryotic cytosol proceeds through rounds of release of native and nonnative forms. Cell 89, 927–937 (1997).

51. R. L. Plimpton et al., Structures of the Gbeta-CCT and PhLP1-Gbeta-CCT complexes reveal a mechanism for G-protein beta-subunit folding and Gbetagamma dimer assembly. Proceedings of the National Academy of Sciences of the United States of America 112, 2413–2418 (2015).

52. S. H. Roh, M. M. Kasembeli, J. G. Galaz-Montoya, W. Chiu, D. J. Tweardy, Chaperonin TRiC/CCT Recognizes Fusion Oncoprotein AML1-ETO through Subunit-Specific Interactions. Biophysical journal 110, 2377–2385 (2016).

53. J. Cuellar et al., Structural and functional analysis of the role of the chaperonin CCT in mTOR complex assembly. Nature communications 10, 2865 (2019).

54. J. Cuellar et al., The structure of CCT-Hsc70 NBD suggests a mechanism for Hsp70 delivery of substrates to the chaperonin. Nat Struct Mol Biol 15, 858–864 (2008).

55. J. Martin-Benito et al., Structure of the complex between the cytosolic chaperonin CCT and phosducin-like protein. Proceedings of the National Academy of Sciences of the United States of America 101, 17410–17415 (2004).

56. J. Martin-Benito et al., Structure of eukaryotic prefoldin and of its complexes with unfolded actin and the cytosolic chaperonin CCT. The EMBO journal 21, 6377–6386 (2002).

57. M. Jin, Y. Cong, Identification of an allosteric network that influences assembly and function of group II chaperonins. Nat Struct Mol Biol 24, 683–684 (2017).

58. T. Lopez, K. Dalton, A. Tomlinson, V. Pande, J. Frydman, An information theoretic framework reveals a tunable allosteric network in group II chaperonins. Nat Struct Mol Biol 24, 726–733 (2017).

59. S. Lu et al., Mapping native disulfide bonds at a proteome scale. Nature methods 12, 329–331 (2015).

60. D. N. Mastronarde, Automated electron microscope tomography using robust prediction of specimen movements. Journal of structural biology 152, 36–51 (2005).

61. S. Q. Zheng et al., MotionCor2: anisotropic correction of beam-induced motion for improved cryo-electron microscopy. Nature methods 14, 331–332 (2017).

62. A. Rohou, N. Grigorieff, CTFFIND4: Fast and accurate defocus estimation from electron micrographs. Journal of structural biology 192, 216–221 (2015).

63. J. Zivanov et al., New tools for automated high-resolution cryo-EM structure determination in RELION-3. Elife 7, (2018).

64. Z. Yang et al., UCSF Chimera, MODELLER, and IMP: an integrated modeling system. Journal of structural biology 179, 269–278 (2012).

65. P. D. Adams et al., PHENIX: a comprehensive Python-based system for macromolecular structure solution. Acta Crystallogr D Biol Crystallogr 66, 213–221 (2010).

66. P. Emsley, K. Cowtan, Coot: model-building tools for molecular graphics. Acta Crystallogr D Biol Crystallogr 60, 2126–2132 (2004).

67. F. DiMaio et al., Atomic-accuracy models from 4.5-A cryo-electron microscopy data with density-guided iterative local refinement. Nature methods 12, 361–365 (2015).

68. J. Jumper et al., Highly accurate protein structure prediction with AlphaFold. Nature 596, 583–589 (2021).

69. J. G. Norby, Coupled assay of Na+,K+-ATPase activity. Methods Enzymol 156, 116–119 (1988).

